# A genetically-defined population of amygdalofugal neurons promotes suckling and early postnatal growth

**DOI:** 10.1101/2025.10.18.683193

**Authors:** Jeffrey D. Moore, Lukas C. Bachmann, Lauren E. McElvain, Samuel L. Pfaff, Catherine Dulac

## Abstract

Suckling by newborns is an instinctive behavior defining the mammalian class. Yet, due to experimental difficulty in assessing neural function in the very young, little is known about the neural control of this fundamental behavior. Here we develop molecular-genetic approaches to interrogate neuronal connectivity and function in newborn mice and used these tools to identify a population of pro-dynorphin (PDYN) and somatostatin (SST) expressing neurons in the central amygdala that are activated during suckling. CeA^PDYN+SST+^ neurons connect with brainstem areas mediating oral sensorimotor and reward function in adults, and their ablation in newborns decreases suckling vigor and impairs growth. These results uncover the crucial role of a specific neuronal population of the central amygdala in maintaining the infants’ propensity to suckle and thrive throughout infancy.

## INTRODUCTION

Suckling behavior is a remarkable ingestive and social adaptation of mammals that enables infants to transiently use maternal resources for nourishment as they develop and mature after birth (*1*). Newborn mammals execute a complex, sensory-guided suckling action sequence that is essential for survival. Additionally, suckling and milk ingestion have been shown to have important anxiolytic and analgesic effects for the infant, including in humans (*2–5*). The suckling sequence involves newborns locating the nipple, attaching to it, and rhythmically sucking to withdraw milk (*6*). Remarkably, suckling is only exhibited in the early postnatal period, and is gradually replaced by independent feeding as the animal matures (*7, 8*). Behavioral studies in laboratory rats demonstrated that nipple attachment is under olfactory and tactile guidance (*9, 10*) and is initially independent of nutritional state (*11–14*). Subsequent studies identified the significant roles of continued suckling experience (*15*), olfactory learning (*16, 17*), behavioral conditioning (*18–21*), and opioid peptides (*22, 23*) to promote and maintain the infant’s propensity to suckle.

However, despite the life-sustaining nature of suckling behavior for mammals, how the central nervous system controls this process is poorly understood. Genetic manipulations to systemically ablate agouti-related peptide-expressing (*AgRP*+) neurons, a neuronal population required for feeding in adult mice, do not appear to affect neonates (*24*), and *AgRP*+ neuronal axon projections do not reach their postsynaptic targets in other hypothalamic brain areas until second postnatal week (*25*). These results suggest that neural circuits that control feeding in mature mammals may not be necessary for infant nutrient ingestion (*26*). However, more precise neural circuit-level investigations at early postnatal stages have been hampered by technical challenges precluding the use of modern neuroscience techniques in infants that are standard in adult rodents. These challenges include the short duration of infancy in rodents; the pliability and rapid changes to the brain and cranium postnatally; and the limited ability to train infant rodents to perform laboratory tasks. Here we develop and employ state of the art molecular-genetic approaches to interrogate the neural architectures that enable the infant mammal to exert behavioral flexibility and control over suckling. We identify a discrete population of prodynorphin (*Pdyn*)*-* and somatostatin (*Sst*)*-*expressing neurons in the central amygdala (CeA), a forebrain structure prominently implicated in emotional learning and motivation (*27–29*), that is specifically active during suckling. We show that CeA^PDYN+SST+^ neurons are required for normal suckling behavior and, in turn, ensuring that the infant grows and thrives.

## RESULTS

### Ontogeny and behavioral flexibility of suckling

As an initial step to identify the central mechanisms controlling suckling behavior, we sought to determine whether and how the suckling motor sequence (*6*) differs according to internal state. Previous studies in rats suggested that the infant’s propensity (probability and latency) to attach to the nipple is independent of internal hunger and satiety cues until the second postnatal week (*11, 12*), prompting us to assess these results in mice (**Fig 1a-d**). We find that mouse pups isolated from their mother for 6 hours on postnatal day 1 (P1) exhibit a decreased latency to attach to the mother’s nipple compared to non-isolated pups (Hazard ratio 4.83 [95% c.i. 2.77-8.44, p=2.9e-8]) (**Fig 1a,b**). This increased propensity to attach could be due to a state-dependent social drive (*30*), suckling drive, or the lack of satiety cues from receiving milk. To distinguish between these different scenarios, we delivered milk intra-gastrically to maternally-deprived pups (**Fig 1c**). Contrary to previous findings in rats (*12–14*), P1 mouse pups who were sham fed exhibited a decreased attachment latency compared to intra-gastrically fed pups (Hazard ratio 7.96 [95% c.i. 2.10-30.15], p=0.002) (**Fig 1d**). These results suggest that, in mice, infant nipple attachment behavior is modulated by internal satiety cues almost from birth.

**Figure 1.**
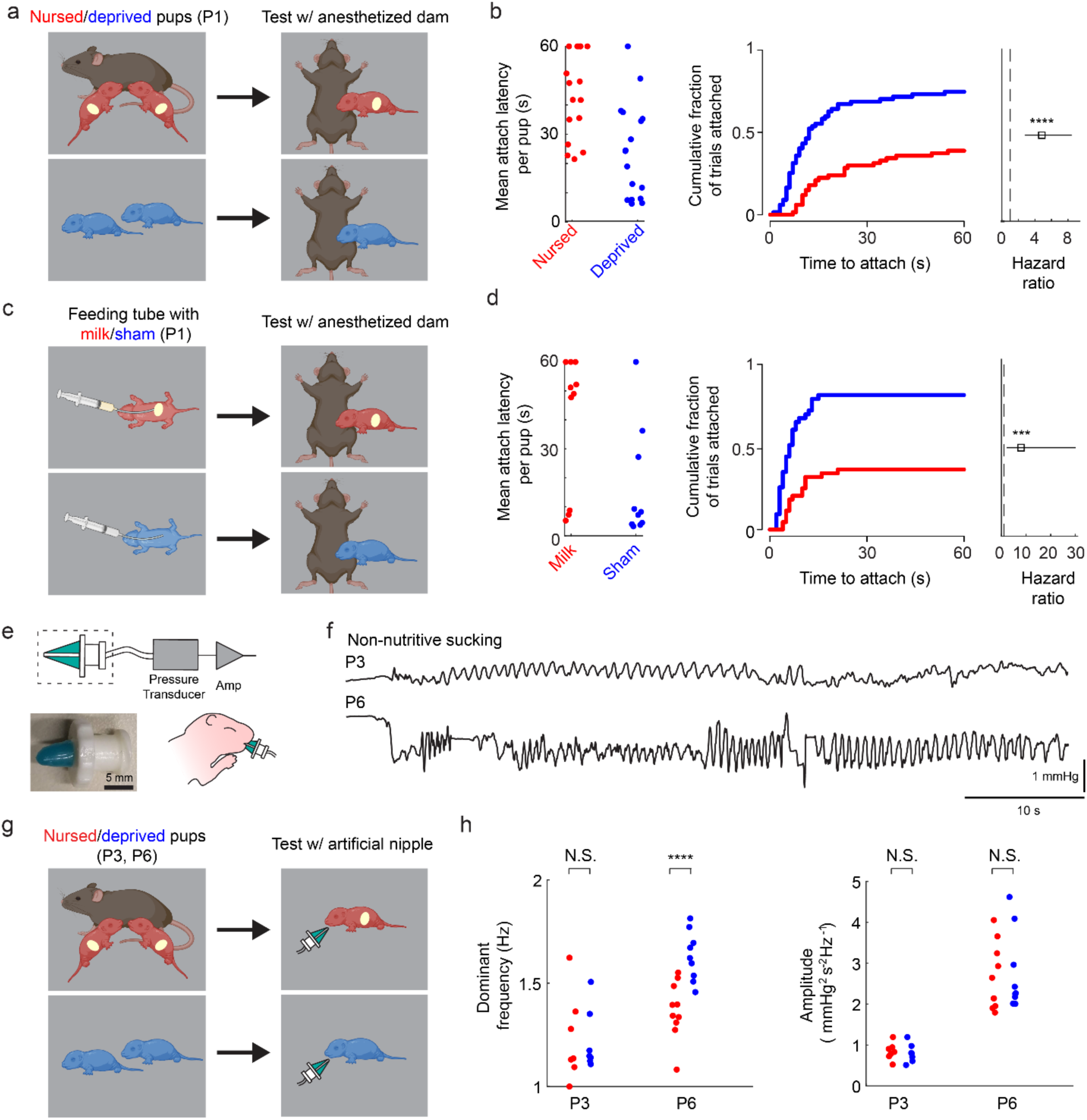
Suckling initiation and dynamics depend on motivational state. **(a)** Experimental paradigm for assessing the effects of maternal separation on attachment to the anesthetized mother. Pups were either left with the mother (red) or isolated (blue) for 6 hours on postnatal day 1 (P1), then manually held adjacent to her ventrum to assess the latency to attach to the nipple. **(b)** (left) Mean latency to attach across all trials for each nursed (red dots) and deprived (blue dots) pup for the experiment in **panel a**. Pups that failed to attach within 60s were assigned a latency of 60s. (middle) Cumulative incidence curves showing the cumulative fraction of trials in which deprived (blue) and nursed (red) pups have attached at each latency for the experiment in **panel a**. (right) Hazard ratio (open square) and 95% confidence interval (line) for the time to attach for deprived versus nursed pups (****p<0.001, Multilevel mixed effects Cox regression; 17 deprived pups and 17 nursed pups in 4 litters) for the experiment in **panel a**. **(c)** Experimental paradigm for assessing the effects of artificial feeding on the attachment to the anesthetized mother. Pups were isolated for 6 hours on postnatal day 1 (P1), then gavage fed with Goat’s milk esbilac (red) or sham fed (blue). They were then manually held adjacent to the anesthetized mother’s ventrum to assess the latency to attach to the nipple. **(d)** (left) Mean latency to attach, (middle) cumulative incidence curves, and (right) Hazard ratio (open square) and 95% confidence interval (line) for the time to attach for sham versus fed pups (****p<0.005, Multilevel mixed effects Cox regression; 10 deprived pups and 10 fed pups in 3 litters). Conventions are as in **panel b (e)** (top) Diagram of an artificial nipple device for assessing reflexive, non-nutritive suckling dynamics in mouse pups. (bottom left) Image of the nipple portion of the device depicted in the dashed box in the diagram. (bottom right) Diagram of the artificial nipple assay in which the pup is held manually and the nipple is coated in oil and inserted into the mouth on P3 or P6. **(f)** Representative recordings of intraoral pressure on P3 and P6. **(g)** Experimental paradigm for assessing the effects of maternal separation on suckling dynamics. Pups were either left with the mother (red) or isolated (blue) for six hours on P3 or P6, and then tested on the artificial nipple assay. **(h)** (left) Mean dominant frequency of rhythmic sucking averaged across all trials per pup at P3 (N.S. – not significant; linear multi-level mixed-effects regression; 7 deprived and 7 nursed pups from 2 litters) and P6 (****p<0.001 at P6 ; 9 deprived and 10 nursed pups from 3 litters). (see Methods). (right) Peak amplitude of rhythmic sucking averaged across all trials for each pup (see Methods).

Studies addressing motivated behavior across a range of species have found that animals tend to modulate the “vigor” of a movement -- defined as its inverse latency (*31*), frequency, or amplitude-- according to its expected utility (*32*). That is, they move with greater speed and lower latency when the movement is likely to yield a valued outcome (*33, 34*). Here, the decreased latency to attach suggests that even at P1, infants may value obtaining milk more when they are deprived than when nursed. We therefore asked whether other aspects of suckling movements are dependent on nutritional state. Specifically, the frequency and intraoral pressure (amplitude) of rhythmic sucking (*7, 12*) are important parameters of suckling that reflect behavioral vigor. To characterize the dynamics of rhythmic sucking, we developed an artificial nipple that was inserted into the pup’s mouth to elicit reflexive, non-nutritive sucking (*35*) while measuring intraoral pressure (**Fig 1e, f**). We find that sucking frequency and amplitude are similar in maternally deprived vs nursed pups at P3 (linear mixed-effects regression, p=0.83 and p=0.74, respectively); (**Fig 1g, h**) however, at P6, suckling-deprived pups suck with a significantly higher frequency than non-deprived littermates (frequency difference = 19.9%, p = 1.7e-7), while amplitude is unaffected (p=0.76) (**Fig 1g, h**). The results demonstrate that, within the first postnatal week, mouse pups can modulate their propensity to attach to the nipple as well as the frequency of rhythmic sucking based on their internal state. Thus, newborn mice appear to have flexible control over suckling vigor at multiple stages of the motor sequence.

### Neuronal populations engaged during suckling

What neuronal circuits enable flexible control of suckling behavior? To address this question, we used an immediate-early-gene screen to broadly survey brain areas whose activity changes when animals suckle or experience sensory and social contexts associated with suckling. P2-P3 mouse pups were isolated for 6 hours and then subjected to one of six behavioral conditions: (1) suckling with their mother; (2) huddling with a non-lactating, experienced mother; (3) continued isolation; (4) periodically agitated or awakened; (5) fed milk intra-gastrically; or (6) stimulated with milk intraorally (**Fig 2a**). Following each behavior, pups were sacrificed and their brains processed for FOS immunolabeling **(Fig S1a-c, Fig 2b)**. To identify brain regions whose activity changes in the holistic behavioral context associated with suckling, we compared FOS immunolabeling in pups who suckled versus those who underwent continued isolation. A number of brain regions exhibited increased FOS expression in suckling versus isolated pups, including the nucleus of the solitary tract (*36*) (NST), the lateral hypothalamus (LH), and the CeA (*37*). We next sought to determine whether other suckling-related activities and stimuli also elicited FOS activation in each region. The NST was also activated after intragastric milk infusion, suggesting a potential role in post-ingestive signaling. The LH was activated following huddling with a non-lactating dam as well as agitation, indicating a possible role in waking or behavioral arousal (**Fig S1a,b**). Among regions that had noticeable FOS expression following suckling, we compared the density of FOS-expressing (FOS+) cells between suckling and huddling conditions (**Fig 2b; Fig S1c**). The CeA appeared as a clear outlier, with far more FOS immunolabeling following suckling than huddling. Specifically, FOS+ neurons in the medial division of the CeA (CeM) were only active when pups were actively suckling compared to all other conditions (**Fig 2c,d**). FOS+ neurons were clustered in the ventrolateral corner of the CeM, and neurons in this same region expressed FOS at P6 and P11 following suckling but not in isolated pups, suggesting suckling-associated activity throughout the early postnatal period (**Fig S1d,e**).

**Figure 2.**
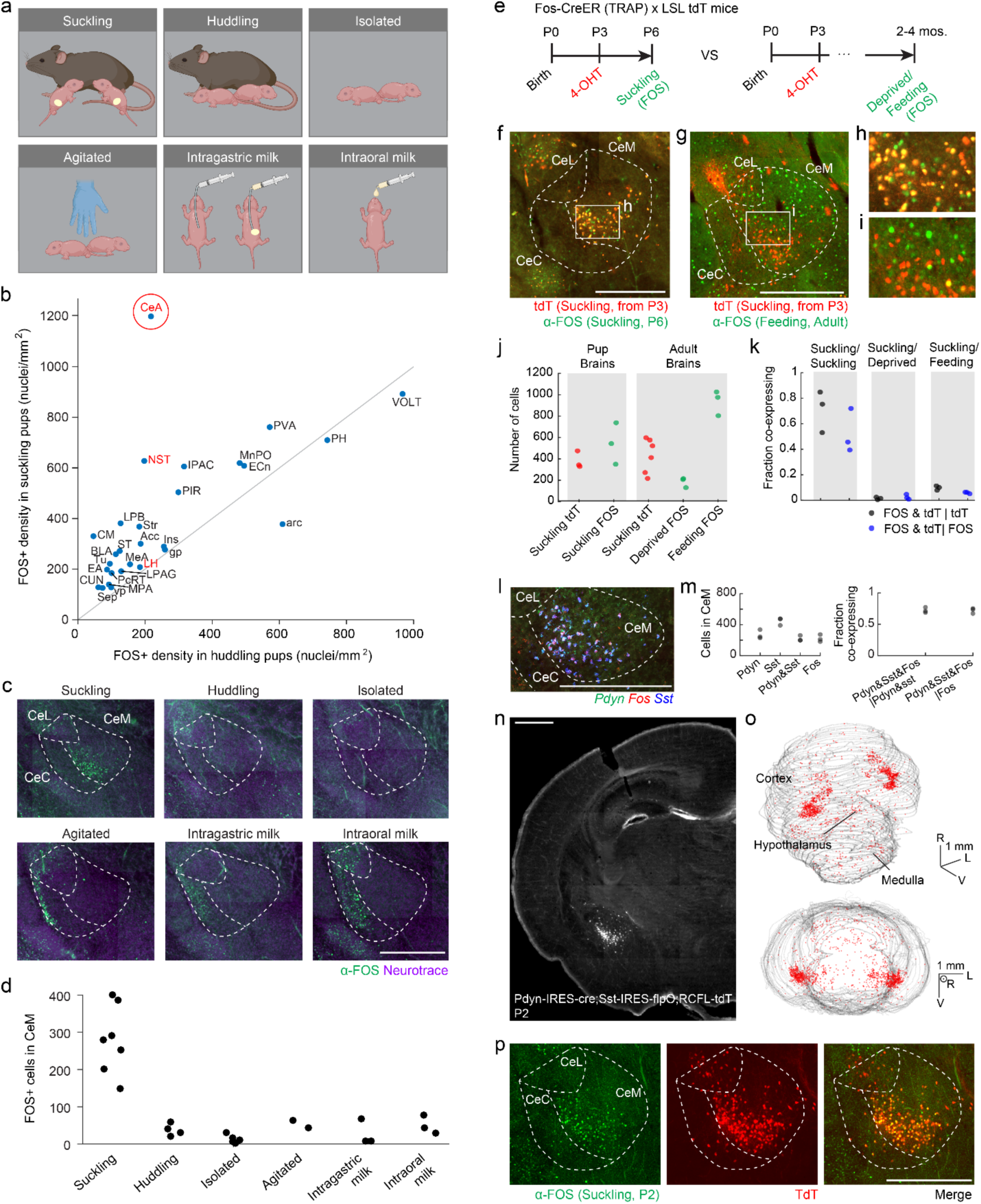
Suckling induces FOS expression in the ventrolateral part of the medial central amygdala. **(a)** Schematics of suckling FOS induction conditions. **(b)** Mean density of FOS immunolabeling in brain regions that were activated in suckling pups (N=4) compared to pups huddling with a non-lactating maternally-experienced, non-lactating female (N=4). Unity line is shown in gray. Example regions that were investigated and depicted in **Fig S1a,b** based on strong differences between suckling and isolated pups are labeled in red. CeA (circled in red) had the greatest difference in density between suckling and huddling conditions. **(c)** Representative images of FOS immunolabeling in the CeA following each of the behavioral conditions in **panel a**. **(d)** Quantification of the number of FOS+ cells in CeM across all behavioral conditions **(e)** Timeline of Fos-CreER induction and subsequent FOS antibody staining. (left) 4-OHT is injected on P3 and suckling assay performed on P6. (right) 4-OHT is injected on P3 and feeding or food deprivation assay is performed in adulthood. **(f)** Representative image of TRAP2;Ai14 TRAPed tdTomato fluorescence (red) and anti-FOS immunolabeling (green) after suckling at P3 and P6, respectively **(g)** Representative image of TRAP2;Ai14 TRAPed tdTomato fluorescence (red) at P3 and anti-FOS immunolabeling (green) after consuming nutrical in adulthood **(h)** blowup of the boxed region in **panel f (i)** blowup of the boxed region in **panel g (j)** (left) Number of TRAPed cells on P3 (red) and FOS+ cells following suckling on p6 (green) (N=3). (right) Number of TRAPed cells on P3 (red) and FOS+ cells (green) following either food deprivation (N=3) or consumption of nutrical (N=3) in adulthood. **(k)** Proportion of cells co-expressing tdTomato and FOS following each of the behavioral paradigms in **panel j**. **(l)** Image of RNAscope (in-situ hybridization) labeling of *Fos* (red), *Pdyn* (green), and *SST* (blue) mRNA in the CeA following suckling at P2. **(m)** (left) Counts of cells in CeM labeled with each of the probes in **panel a.** (right) proportion of CeM *Pdyn+,Sst+* cells that also express *Fos* (black) and proportion of CeM *Fos*+ (suckling) cells that are also *Pdyn+,Sst+* (blue) (N=3) **(n)** representative section through the CeA of a Pdyn-IRES-cre(+/-);Sst-IRES-flpO(+/-);Ai65(+/-) (RCFL-tdT) mouse at P2 **(o)** (top) Whole brain reconstruction of tdTomato+ cells for the mouse in **panel n**, oblique view. (bottom) Frontal view of the same brain reconstruction. Rostral (R), lateral (L), ventral (V) directions are indicated. **(p)** Representative image of a P2 Pdyn-IRES-cre(+/-);Sst-IRES-flpO(+/-);Ai65(+/-) mouse with: (left) anti-FOS immunolabeling following suckling, (middle) tdTomato expression in the same field of view, (right) an overlay of these two images. Scale bars are 500um in all panels unless otherwise indicated. Anatomical abbreviations: Central amygdala (CeA), Nucleus of the solitary tract (NST), interstitial nucleus of the posterior limb of the anterior commissure (IPAC), centromedian thalamuc (CM), lateral parabrachial area (LPB), piriform cortex (PIR), anterior nucleus of the paraventricular thalamus (PVA), striatum (Str), bed nuceus of the stria terminalis (ST), basolateral amygdala (BLA), median preoptic nucleus (MnPO), olfactory tubricle (Tu), nucleus accumbens (Acc), external cuneate (ECn), extended amygdala (EA), parvocellular reticular formation (PcRT), cuneate nucleus (CUN), medial amygdala (MeA), lateral periacqueductal gray (LPAG), septum (Sep), medial preoptic area (MPA), insular cortex (ins), ventral pallidum (vp), lateral hypothalamus (LH), globus pallidus (gp), posterior hypothalamus (PH), vascular organ of the lamina terminalis (VOLT), arcuate nucleus (arc), medial division of the central amygdala (CeM), lateral division of the central amygdala (CeL), capsular division of the central amygdala (CeC).

The CeA has been implicated in adult appetitive behaviors including palatable food and fluid consumption and reward learning (*29, 38–40*). We therefore asked whether the same CeA neurons are associated with both feeding in adults and suckling in newborns. We used the TRAP2 mouse line, which expresses tamoxifen-inducible creER under the *Fos* promoter (2A-iCreER^T2^) (*41*) to permanently label neurons that are active during newborn suckling (**Fig 2e-k)**. Specifically, TRAP2;Ai14 (Lox-Stop-Lox (LSL) tdTomato) (*42*) mouse pups were injected with 4 hydroxy-tamoxifen (4-OHT) at P2-3 to induce cre-dependent tdTomato expression. To determine the extent to which this paradigm captures suckling active neurons in the CeA, the same mice were subjected to maternal separation followed by nursing at P6, and subsequently sacrificed (**Fig 2e**). Brains were then processed for FOS immunolabeling. Within CeA, tdTomato fluorescence and FOS immunolabeling were largely restricted to the ventrolateral CeM (**Fig 2f,h**). A high fraction of the tdTomato+ neurons induced at P2-3 were also FOS+ at P6 (72% [95% c.i.: 56-85%]) and, likewise, a high fraction of FOS+ neurons at P6 were also tdTomato+ from TRAP at P2-3 (53% [95% c.i. 36-69%]) (**Fig 2j,k**). This observation suggests that both TRAP2-labeling and direct FOS immunolabeling identify a largely similar population of suckling active neurons at P3 and P6, respectively.

Next, TRAP2;Ai14 mouse pups induced with 4-OHT on P2-3 were allowed to grow to adulthood and undergo a feeding assay (**Fig 2e**). In this assay, adult mice were either presented with palatable food (nutrical) or kept deprived. Mice were then sacrificed and brains were processed for FOS immunolabeling (**Fig 2g,i**). The brains of fed mice displayed an approximately 5-fold increase in the number of FOS+ cells in CeM compared to deprived mice (p=4.8 x 10^-4, t-test) (**Fig 2j**). By contrast, mice who were allowed to drink palatable fluid (2% sucrose) did not show appreciable FOS labeling in CeM (**S1f**). Strikingly, neurons activated following feeding were distributed throughout the CeM and were largely distinct from the population of suckling-active neurons. That is, only 5.9% [95% c.i.: 5.1-6.9%] of TRAPed cells on P2-3 were also FOS+ after feeding, and 9.8% [95% c.i.: 8.5-11.3%] of Fos+ cells were also TRAPed (**Fig 2k**). These observations suggest that the limbic circuits involved in infant suckling are distinct from those associated with adult ingestive behaviors (*29*).

### Cell-types of suckling-active CeA neurons

The CeA has been shown to contain transcriptionally diverse neuronal populations expressing a variety of neuropeptides (*29, 43*). To identify molecular markers of suckling-active CeM neurons, we performed in-situ hybridization experiments in P2-3 mice to co-label *Fos* mRNA following suckling along with candidate genes known to be expressed in the CeA (*29*), including *pro-dynorphin (Pdyn), somatostatin (Sst), neurotensin (Nts), vesicular GABA transporter (vGat), prepronociceptin (Pnoc),* and *Delta-like non-canonical notch ligand 1 (Dlk1)*(*29, 44–46*) (**Fig 2l, Fig S2a-c**). As with FOS protein, *Fos* mRNA was increased following suckling compared to deprived littermates (**Fig S2a**). Moreover sucking-induced *Fos+* neurons in CeM significantly co-localized with expression of *Pdyn* and *Sst*, with a high fraction of cells expressing both neuropeptides (**Fig 2l,m, Fig S2a,c**). Specifically, 63% [95% c.i. 60-67%] of *Pdyn* cells expressed *Fos*, and 42% [95% c.i. 37-46%] of *Sst* cells expressed *Fos.* Of cells that expressed both *Pdyn* and *Sst*, 73% [95% c.i. 69-76%] expressed *Fos*, and of cells that expressed *Fos*, 72% [95% c.i. 68-76%] expressed both *Pdyn* and *Sst.* All of the cells expressing *Pdyn* and/or *Sst* also express *vGat*, as expected from observations that the CeA is overwhelmingly GABAergic (*43, 47*) (**Fig S2d,e**). These findings suggest that within the CeA, suckling specifically activates inhibitory neurons characterized by the *co-*expression of *Pdyn* and *Sst*.

### Cell-type-specific genetic access to CeA neurons active during suckling

The molecular identification of specific CeM neurons active during suckling provided genetic entry points to characterize their connectivity with other brain regions and assess whether and how these neurons may drive or support behavior. Gaining spatially-restricted and cell-type-specific genetic access to neuronal populations is typically achieved by the stereotaxic injection of conditional viral vectors expressing reporters or actuators into discrete brain regions of transgenic mouse lines, most commonly with adeno-associated viral (AAV) vectors (*48, 49*). However, achieving such spatial and genetic specificity during early postnatal development presents several technical challenges. In part, the brain and cranium are smaller and more pliable than in adult mice, presenting a physical challenge as it hinders precise spatial targeting during stereotaxic surgery.

To assess genetic access to the specific CeA neuronal population activated during suckling, we examined mouse lines that drive cre-recombinase in cells that express either *Pdyn* or *Sst*, crossed to a cre-dependent tdTomato reporter (LSL-tdT, Ai14) mouse at early postnatal ages (Fig **S2f-j**). Both Pdyn-IRES-cre;Ai14 (*50*) and Sst-IRES-cre;Ai14 mice (*51*) exhibit a large number of tdTomato+ neurons in the CeA as well as in nearby brain regions, making it challenging to target the CeA population specifically by viral injection in newborn mice (**Fig S2i, j**). Additionally, in Sst-IRES-cre;Ai14 mice, tdTomato expression within CeA did not appear to be spatially restricted to the ventrolateral CeM (**Fig 2c** versus **Fig S2f-h**). We next checked whether intersectional crosses of Pdyn-IRES-cre and Sst-IRES-flpO mice would result in greater genetic and anatomical specificity. We generated Pdyn-IRES-cre(+/+);Sst-IRES-flpO(+/+) mice and crossed them to a homozygous cre- and flp-dependent tdTomato reporter mouse line (RCFL-tdT, Ai65) (*52, 53*), resulting in Pdyn-IRES-cre(+/-);Sst-IRES-flpO(+/-);Ai65(+/-) pups, whose brains we examined histologically. These mice exhibit tdTomato+ neurons in the CeA (CeA^PDYN+SST+^ neurons) and, importantly, relatively few neurons in surrounding brain areas (**Fig 2n,o**). Within CeM, 59% [95% c.i. 50-67%] of these tdTomato+ neurons co-express FOS following suckling, and 70% [95% c.i. 68-71%] of FOS+ cells are tdTomato+ (**Fig 2p, Fig S2k**) Across all CeA subdivisions, 52% [95% c.i. 43-61%] of tdTomato+ neurons are FOS+ following suckling, and 48% [95% c.i. 45-50%] of FOS+ cells are tdTomato+. We estimate 16% of NeuN+ cells in CeM and 12% of NeuN+ neurons in all CeA subdivisions are tdTomato+, corresponding to 4.4-fold and 4-fold enrichment ratios for FOS (fraction of FOS+ co-expressing tdTomato divided by the fraction of NeuN+ cells expressing tdTomato) (*54*), in CeM and CeA, respectively. We therefore sought to use the Pdyn-IRES-cre(+/+);Sst-IRES-flpO(+/+) intersectional mouse line to characterize the input and output projections of CeA^PDYN+SST+^ neurons, and whether and how they may regulate suckling behavior.

### Development of an scAAV-based strategy for rapid axonal tracing

It is well-established that postnatal axonal projections evolve over time with sequential maturation steps at different postnatal ages (*25*). In mice, the short early-postnatal period (3 weeks from birth until weaning) creates a timing challenge because axonal tracing from neurons requires significantly high levels of viral transgene expression that typically take 2-4 weeks with most commercially available AAV vectors (*55*). Therefore, identifying axonal projections and axon terminal locations that are present at early ages requires techniques for rapid, cell-type specific axonal tracing techniques. To address this issue, we developed a self-complementary adeno-associated viral vector (scAAV) and an experimental strategy to gain rapid genetic access to neurons and trace their anatomical connectivity. scAAVs have been demonstrated to exhibit markedly faster expression times and higher expression levels than standard single-stranded AAV (ssAAV) vectors, albeit with smaller transgene payload sizes (*55, 56*). To map the projection targets of specific neurons that are present in early life, we developed a cre-dependent scAAV vector expressing a molecularly tagged synaptophysin protein (scAAV-hSyn-FLEx-Synaptophysin-HA) to map pre-synaptic axon terminals, and used it in conjunction with a similar rapid GFP expression vector targeting somata and axons (scAAV-hSyn-FLEx-GFP (*57*)) (**Fig 3a**). These viruses were co-injected into the amygdala of Pdyn-IRES-cre(+/-);Sst-IRES-flpO(+/-);Ai65(+/-) pups at P1 (**Fig 3b**). After 7 days, pups were sacrificed and brains were processed immunohistochemically with anti-GFP, anti-RFP, and anti-HA antibodies. We observed GFP labeling of putative *Pdyn*-expressing neurons in and around the amygdala (**Fig 3c**). Somata of CeA^PDYN+SST+^ neurons as well as their axons are double-labeled with GFP and tdTomato (**Fig 3c**, yellow), and presynaptic boutons arising from these neurons additionally expressed the synaptophysin-HA tag in the caudal-most regions of the medulla (**Fig 3d**, yellow), suggesting that a one week incubation period is sufficient to label axonal projections and terminals from CeA neurons throughout the entire brain.

**Figure 3.**
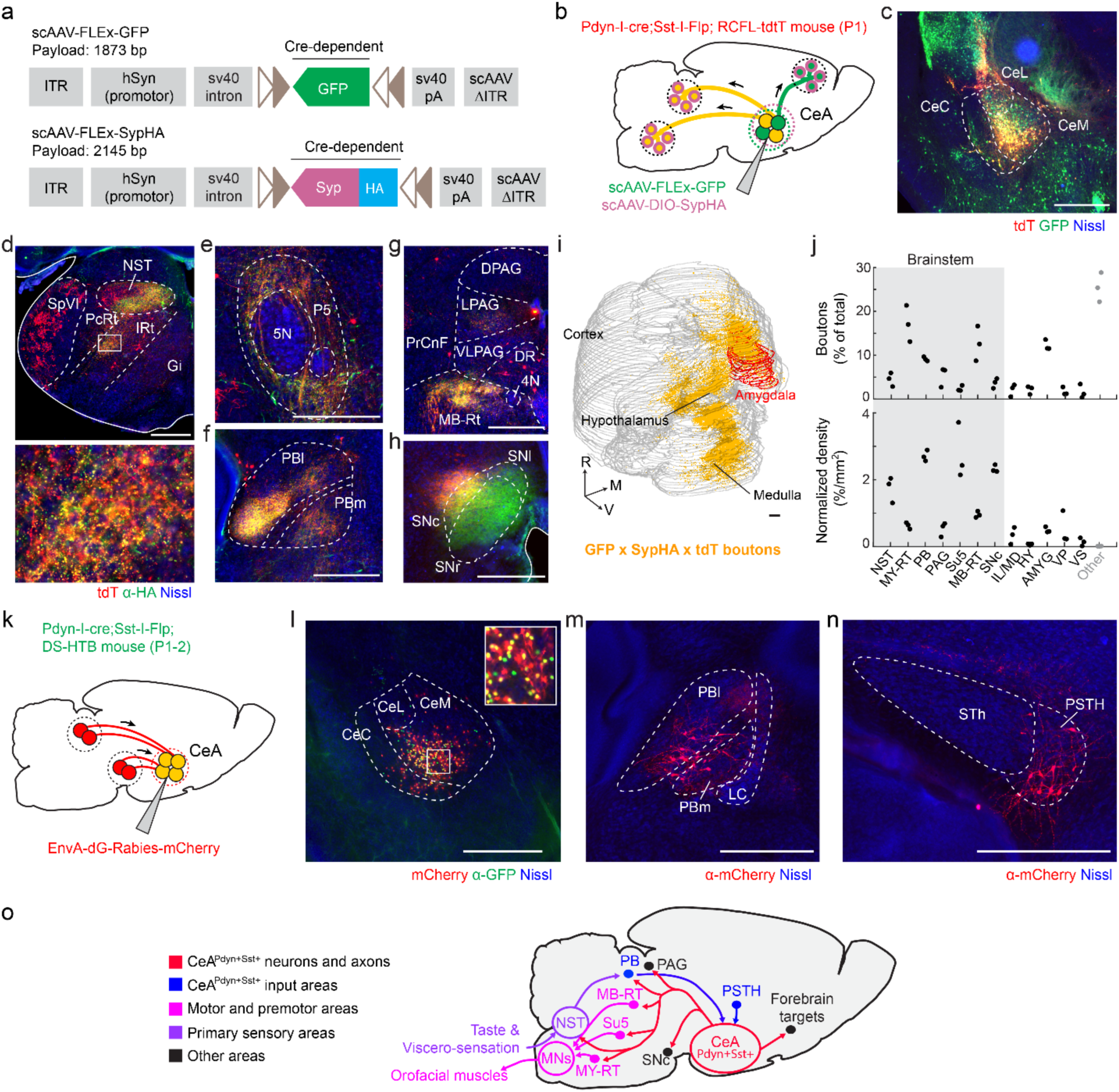
Brain-wide connectivity of CeA^PDYN+SST+^ neurons at early postnatal ages is consistent with a role in influencing suckling behavior. **(a)** Map of scAAV vectors for rapid tracing of axons and axon terminals in cre-expressing neurons. Abbreviations: self-complementing adeno-associated virus (scAAV), ITR (inverted terminal repeat), polyadenylation tail (pA), green fluorescent protein (GFP), human synapsin (hSyn), synaptophysin (Syp), hemagglutinin tag (HA). The number of base pairs in the construct between the two ITR regions is indicated. *Lox* sites are shown in light brown **(b)** Strategy for labeling outputs of CeA^PDYN+SST+^ neurons. Injections of scAAV2/8-hSyn-FLEx-GFP and scAAV2/8-hSyn-FLEx-SypHA are roughly targeted stereotaxically to the amygdala of Pdyn-IRES-cre(+/-);Sst-IRES-FlpO(+/-);Ai65(+/-) mice on P1. Cre+/Flp-neurons and their axons will be labeled with GFP (green) and their axon terminals will additionally be labeled with HA-tagged synaptophysin (magenta), Cre+/Flp+ neurons and their axons and terminals will be labeled with tdTomato, and Cre+/Flp+ neurons within the injection site and their axons will be labeled with both tdTomato and GFP (yellow) and their axons will additionally be labeled with HA (magenta). The mice are sacrificed at P8. **(c)** Image of anti-RFP and anti-GFP immunolabeling following the paradigm described in **panel b.** Cre+FlpO+ cells co-express GFP (green), tdTomato (red), and HA (not shown) while Cre+FlpO-cells express only GFP and HA, but not tdTomato. **(d)** Brain-wide immunolabeling for anti-HA (green) and anti-tdTomato (red) reveal co-labeled terminals in the NST and IRt/PcRt (top) and blowup of rectangle region (bottom) **(e)** Terminals are also double-labeled in the P5 region, **(f)** PBl and PBm **(g)** MB-Rt and subdivisions of the PAG, and **(h)** SNc. **(i)** Sample whole brain reconstruction of terminals triple-labeled with anti-GFP, anti-tdTomato, and anti-HA (yellow) and the amygdala boundaries (red) shown at an oblique view. The rostral (R), medial (M), ventral (V) directions are indicated. **(j)** Quantification of relative numbers of boutons (top) and bouton density (bottom) in highly innervated brain regions ipsilateral to the injection site (black) and the remaining boutons throughout the brain (gray) **(k)** Strategy for labeling pre-synaptic inputs to CeA^PDYN+SST+^ neurons. Injections of EnvA-SADB19-dG-Rabies-mCherry are roughly targeted to the CeA in Pdyn-IRES-cre(+/-);Sst-IRES-flpO(+/-); Rosa-HTB(+/-) mice at P1-2. The mice are sacrificed on P9 or P14. Starter cells (yellow) express GFP in the nucleus and mCherry in the soma, and neurons that provide presynaptic inputs express mCherry only (red). **(l)** Image from a P9 mouse labeled with anti-GFP showing Cre+FlpO+ cells co-expressing GFP (green) in nuclei and mCherry (red) in somata (putative starter cells), and a blowup of the rectangle region (inset) **(m)** Image of EnvA-SADB19-dG-Rabies-mCherry labeling in PBl and PBm and **(n)** PSTH. **(o)** Summary diagram of identified input-output connectivity of CeA^PDYN+SST+^ neurons in the mouse pup. CeA^Pdyn+Sst+^ neurons (red) get inputs from feeding associated areas (blue) and target their outputs to orofacial premotor structures (magenta), brainstem primary sensory areas (purple), as well as other areas associated with motivated behavior (black). Scale bars are 500um in all panels. Anatomical abbreviations: Cetral amygdala medial, lateral and capsular divisions (CeM, CeL, CeC, respectively), Spinal trigeminal nucleus interpolaris (SpVI), parvocellular reticular formation (PcRT), nucleus of the solitary tract (NST), intermediate reticular formation (IRt), gigantocellular reticular formation (Gi), Trigeminal motor nucleus (5N), peritrigeminal region (P5), parabrachial area (PB) lateral and medial divisions (PBl and PBm, respectively), precuniform (PrCnF), midbrain reticular formation (MB-RT), periacueductal gray (PAG) dorsal, lateral, ventolateral divisions (DPAG, LPAG, VLPAG, respectively), 4^th^ motor nucleus (4N), dorsal raphe (DR), substantia nigra compact, reticular, and lateral divisions (SNc, SNr, SNl respectively), medullary reticular formation (MY-RT), supratrigeminal region (Su5), interlaminal/medial thalamus (IL/M), hypothalamus (HY), amygdala (AMYG), ventral pallidum (VP), ventral striatum (VS), locus coeruleus (LC), subthalamic nucleus (Sth), paratubthalamic nucleus (PSTH).

### Projections of CeA^PDYN+SST+^ neurons

We used this dataset to map which downstream brain areas are innervated by CeA^PDYN+SST+^ neurons during infancy. GFP+tdTomato+ somata were largely restricted to the CeA (**Fig 3c**), and triple-labeled presynaptic boutons from these neurons were identified in a number of long range projection targets including forebrain projections to the ipsilateral ventral striatum (VS), hypothalamus (Hy), and the interlaminar/medial thalamic nuclei (IL/M) (*58*), along with dense projections along the amygdalofugal pathway to discrete brainstem areas including the NST and the medullary parvocellular and intermediate reticular formation (PcRt/IRt) (**Fig 3d**), the peri-trigeminal region (P5) (**Fig 3e**), the parabrachial area (PB) (**Fig 3f**), midbrain reticular formation (MRt) (**Fig 3g**), and the lateral aspect of the substantia nigra pars compacta (SNc) (**Fig 3h**). We digitally reconstructed the triple-labeled boutons arising from CeA^PDYN+SST+^ neurons throughout the brain (**Fig 3i, Fig S3a**). The ipsilateral brainstem outputs of these neurons, which comprise the majority of the total synaptic boutons (**Fig 3j**), were highly specific for regions that contain premotor neurons innervating oro-motor neurons (MRt, peri-V, IRt) in pups (*59*), oro- and viscero-sensory areas (NST, PB), and the midbrain dopamine system (SNc). These projection patterns, which comprise a subset of known output projections of CeA neurons in adult mice (*43, 60, 61*), are consistent with the notion that CeA^PDYN+SST+^ neuronal activity may influence orosensory processing, oromotor function, and/or reward signaling.

### Synaptic inputs to CeA^PDYN+SST+^ neurons

We next sought to identify synaptic inputs to CeA^PDYN+SST+^ neurons using conditional dG-rabies tracing (*62*). To do this, we used a mouse line that enables tracing synaptic inputs from starter cells that express both cre-and flp-recombinase (Rosa-DS-HTB) (*63, 64*). This mouse line expresses nuclear bound GFP (H2B-GFP), the avian ASLV-A receptor (TVA), and rabies glycoprotein (B19G), in the presence of both cre- and flp-recombinases. We crossed Rosa-DS-HTB(+/+) mice to Pdyn-IRES-cre(+/+);Sst-IRES-flpO(+/+) mice, and EnvA-SAB19-dG-rabies-mCherry was injected into the CeA in P1-2 pups (**Fig 3k-l**). Intermingled with cells expressing both GFP and mCherry, the putative CeA^PDYN+SST+^ “starter neurons”, are cells that are exclusively mCherry+, which could represent local presynaptic interneurons or non-specific rabies entry to cells within the injection site (*65*). We also consistently observed mCherry+ neurons in the parabrachial area and parasubthalamic nucleus (**Fig 3m,n; Fig S3b**), which putatively send long-range pre-synaptic inputs to CeA^PDYN+SST+^ neurons. While the low efficiency of this tracing strategy (*66*) makes it likely that these are not the only sources of input, their labeling suggests they represent prominent input sources. As with the output targets, these regions have been shown to be involved in integrative oro- and viscero-sensory functions (*67, 68*). Together the pattern of input-output connectivity of the CeA^PDYN+SST+^ neurons (**Fig 3o**) is consistent with a putative role in regulating suckling behavior.

### New scAAV vectors to rapidly manipulate intersectionally-defined neurons

We next sought to specifically ablate CeA^PDYN+SST+^ neurons in pups to determine whether and how these neurons may influence suckling behavior. We developed an scAAV vector that expresses diphtheria-toxin subunit A (DTA) in a cre- and flp-dependent manner via RNA splicing (*69, 70*), as well as a corresponding GFP-expressing control vector (**Fig 4a,b**). These vectors meet the reduced size constraint required for scAAV packaging (*56*) by using a truncated EF1a promoter (EF1a(core)), unidirectional *lox* and *frt* mutant recombination sites, and modified splice sites (scAAV2/8-EF1a(core)-DDIO-DTA and scAAV2/8-EF1a(core)-DDIO-GFP). We reasoned that, in addition to more rapid expression than ssAAV vectors to gain genetic access to neurons in early life, the increased expression levels of scAAVs may additionally overcome limitations in the efficiency of dual-recombinase splice-based expression strategies (*71*). To test the effectiveness and specificity of this strategy, we injected scAAV2/8-EF1a(core)-DDIO-GFP into the amygdala in Pdyn-IRES-cre(+/-);Sst-IRES-flpO(+/-);Ai65(+/-) mouse pups at P1. GFP fluorescence was detected after as little as 5 days, and the labeling was restricted to CeA tdTomato+ neurons (**Fig S4a**). At P14, there was more variability in tdTomato fluorescence levels within GFP+ cells (**Fig 4c**); specifically, 77% (95% c.i. 75-79%, N=3) of GFP+ neurons had detectable tdTomato fluorescence. Nonetheless, GFP fluorescence was not observed outside the CeA sub-region containing Pdyn+,Sst+ neurons (**Fig 4c**, **Fig 2l**), suggesting that off-target expression of GFP is unlikely. Furthermore, injecting scAAV2/8-EF1a(core)-DDIO-GFP into the amygdala in Pdyn-IRES-cre(+/-);Sst-IRES-flpO(-/-);Ai65(+/-) littermates resulted in no off-target labeling (**Fig S4b**). Together, these observations demonstrate that the scAAV2/8-EF1a(core)-DDIO construct is able to drive rapid and specific expression in CeA^PDYN+SST+^ neurons.

**Figure 4.**
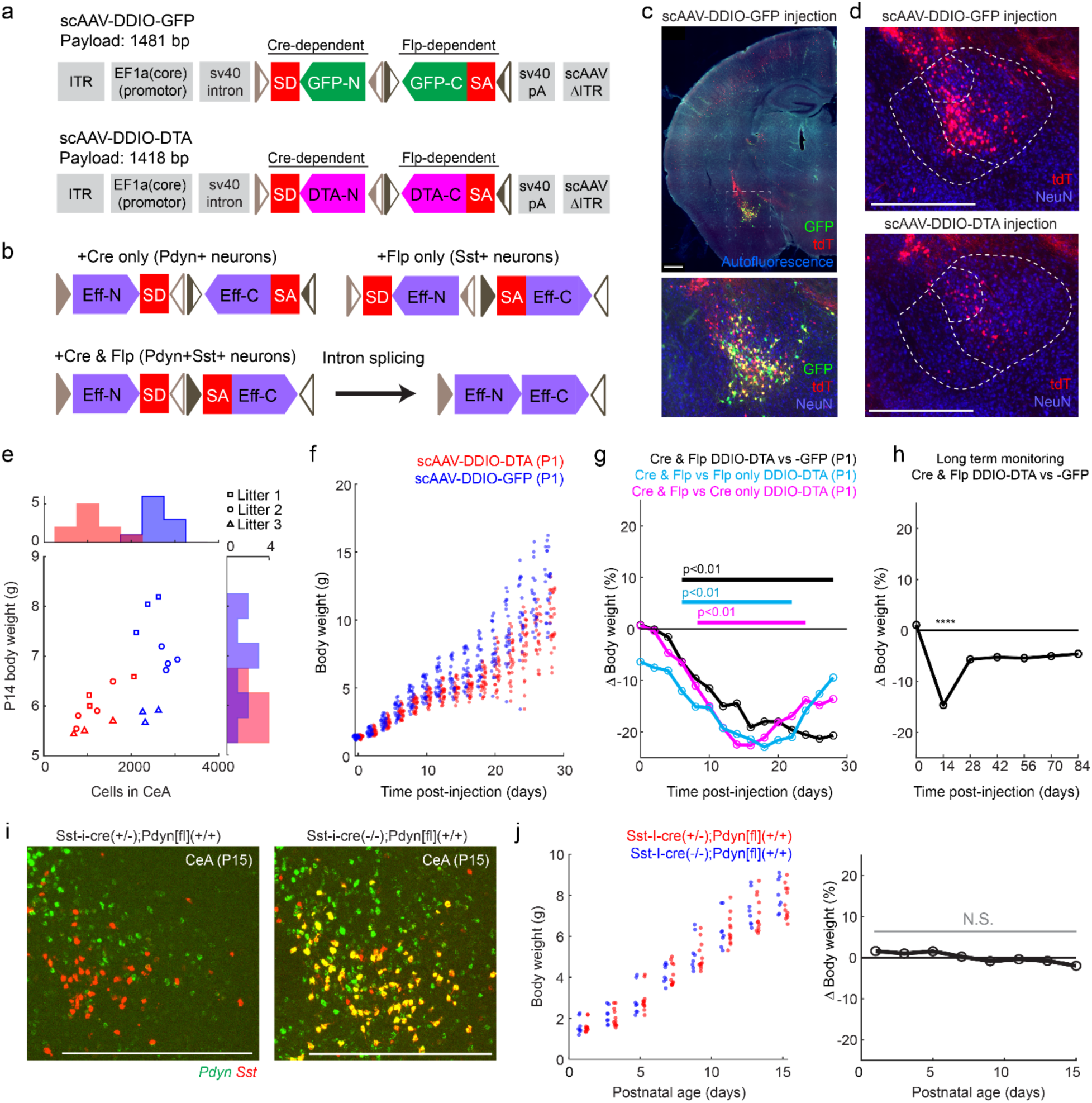
Rapid viral ablation of CeA^PDYN+SST+^ neurons leads to growth deficits in early life. **(a)** Design of cre- and flp-dependent scAAV vectors expressing GFP (scAAV-EF1a(core)-DDIO-GFP) (top) and diphtheria toxin subunit A (DTA) (scAAV-EF1a(core)-DDIO-DTA) (bottom). Conventions are as in Figure 3a. Abbreviations: splice donor site (SD), splice acceptor site (SA), N-terminus (-N), C-terminus (-C), diphtheria toxin (DTA). Unidirectional *lox* mutant sites are shown in light brown, and unidirectional *frt* mutant sites are shown in dark brown **(b)** Schematic of expected mRNA expression in neurons expressing cre-recombinase only (upper left), flp-recombinase only (upper right), and both recombinases before (lower left) and after (lower right) intronic splicing. The neuronal populations expected to express each construct in Pdyn-IRES-cre;Sst-IRES-FlpO mice is indicated. Abbreviations: Effector gene (Eff) refers to either GFP or DTA in **panel a**. **(c)** (top) Image of a coronal section through the CeA of a P14 Pdyn-IRES-cre(+/-);Sst-IRES-flpO(+/-);Ai65(+/-) mouse with an injection of scAAV2/8-EF1a(core)-DDIO-GFP targeted to the amygdala at P1. (bottom) Blowup of the CeA from the outlined area in the image above with an anti-NeuN counterstain **(d)** (top) tdTomato expression and anti-NeuN immunolabeling in CeA of a Pdyn-IRES-cre(+/-);Sst-IRES-flpO(+/-);Ai65(+/-) mouse following injection of scAAV2/8-EF1a(core)-DDIO-GFP and (bottom) scAAV2/8-EF1a(core)-DDIO-DTA. **(e)** body weight versus number of remaining tdTomato+ cells in CeA for mice injected with scAAV2/8-EF1a(core)-DDIO-DTA (red) or scAAV2/8-EF1a(core)-DDIO-GFP (blue) at P1 and sacrificed at P14. Each symbol type represents pups from a single litter. Histograms of the distributions of remaining cells and body weight are shown for each group along the corresponding axis **(f)** Body weights measured every two days from P1 to P28 following P1 injections of scAAV2/8-EF1a(core)-DDIO-GFP (blue, N=29 from 8 litters) or -DTA (red, N=27 littermates) targeted to the CeA in Pdyn-IRES-cre(+/-);Sst-IRES-flpO(+/-);Ai65(+/-) mouse pups. In each litter, approximately half the pups were injected with each vector. **(g)** Estimates of differences in body weights between ablated mouse pups versus non-ablated littermates for the corresponding experimental conditions in **panel f** and **Fig S4c.** Relative body weights are assessed with a linear mixed-effects regression model that accounts for litter effects (see Methods). Bars represent statistical significance at the p<0.01 level for each cohort. **(h)** Estimates of differences in body weights between Pdyn-IRES-cre(+/-);Sst-IRES-flpO(+/-);Ai65(+/-) mice with P1 injections of scAAV2/8-EF1a(core)-DDIO-DTA (N=15 from 5 litters) and littermates with P1 injections of scAAV2/8-EF1a(core)-DDIO-GFP (N= 15). These data are from a separate cohort of mice from those in **panel f** in which body weight was measured every two weeks for 10-12 weeks. Relative body weights are assessed with a linear mixed-effects regression model that accounts for litter effects. **** p<0.0001. **(i)** Hybridization chair reaction (HCR®) in-situ hybridization for *Sst* (red) and *Pdyn* (green) in Sst-IRES-cre(+/-);Pdyn[fl](+/+) (left), and Sst-IRES-cre(-/-);Pdyn[fl](+/+) (right) showing the CeA of P15 pups. **(j)** (left) Postnatal body weights of Sst-IRES-cre(+/-);Pdyn[fl](+/+) mouse pups (red, N=12 from 5 litters) and Sst-IRES-cre(-/-);Pdyn[fl](+/+) littermates (blue, N=8 from 5 litters) resulting from crosses of Sst-IRES-cre(+/-);Sst-IRES-cre(+/-) and Pdyn[fl](+/+) mouse lines. (right) Estimates of differences body weights between the Sst-IRES-cre(+/-);Pdyn[fl](+/-) group and the Sst-IRES-cre(+/-);Pdyn[fl](-/-) littermate group. Relative body weights are assessed with a linear mixed-effects regression model that accounts for litter effects. N.S. – not significant at p<0.05. Scale bars are 500um in all panels.

### CeA^PDYN+SST+^ neurons promote early postnatal growth

We employed this newly developed scAAV-based intersectional viral gene-delivery strategy to specifically ablate the CeA^PDYN+SST+^ neuronal population and assessed behavioral and physiological changes in early postnatal mice. At P1, we injected half of the pups in litters of Pdyn-IRES-cre(+/-);Sst-IRES-flpO(+/-);Ai65(+/-) mice with scAAV2/8-EF1a(core)-DDIO-DTA, and the remaining pups in each litter with scAAV2/8-EF1a(core)-DDIO-GFP. We examined the GFP-labeling (**Fig 4c**) and DTA-ablation efficacy (**Fig 4d**) at P14, and we confirmed a 54% reduction (p=6.8e-8, linear mixed-effects regression model accounting for litter effects) in the number of tdTomato+ cells in DTA-compared to GFP-injected control mice (**Fig 4e**). Moreover, a 14% decrease in body weight (p=2.4e-5, linear mixed-effects regression) was observed at P14 in DTA-ablated pups. Among these pups, body weight significantly correlated with the number of remaining CeA^PDYN+SST+^ neurons at that stage (5.23g + 0.6mg/cell, p=0.0018, linear mixed-effects regression model, N=10 pups) (**Fig 4e**). To quantify the time course of the effects of CeA^PDYN+SST+^ neuronal ablation on body weight, we injected a separate cohort of Pdyn-IRES-cre(+/-);Sst-IRES-flpO(+/-);Ai65(+/-) mice with scAAV2/8-EF1a(core)-DDIO-DTA or -GFP at P1, and tracked body weights from birth until P28. DTA-treated pups exhibited markedly slower growth than GFP-injected littermates with statistically significant (p<0.01) deficits in body weights observed beginning at P6 and accumulating throughout the early postnatal period (**Fig 4f**; **Fig 4g –** black trace).

Since the scAAV2/8-EF1a(core)-DDIO-GFP and scAAV2/8-EF1a(core)-DDIO-DTA viral vectors undergo recombination of a cre-dependent DNA segment in the presence of cre-alone, and recombination of a flp-dependent DNA segment in the presence of flp-alone (**Fig 4a,b**), we sought to verify the specificity of the ablation by checking whether the observed effects on growth may be due to off-target effects of DTA in nearby Pdyn+/Sst-cells or Pdyn-/Sst+ neurons. We crossed Pdyn-IRES-cre(+/-);Sst-IRES-flpO(+/+) mice or Pdyn-IRES-cre(+/+);Sst-IRES-flpO(+/-) mice with Ai65(+/+) mice, resulting in litters comprised of pups that expressed *both* cre- and flpO-recombinase in *Pdyn-* and *Sst-*expressing neurons, as well as littermates expressing *either* cre- or flpO-recombinase in these neurons. We injected scAAV2/8-EF1a(core)-DDIO-DTA in the CeA of all pups in each litter on P1. As in the previous experiment (**Fig 4f**), we observed growth deficits when comparing Pdyn-IRES-cre(+/-);Sst-IRES-flpO(+/-);Ai65(+/-) pups to either Pdyn-IRES-cre(+/-);Sst-IRES-flpO(-/-);Ai65(+/-) or Pdyn-IRES-cre(-/-);Sst-IRES-flpO(+/-);Ai65(+/-) littermates (**Fig S4c, Fig 4g**), suggesting that growth deficits are indeed due to DTA expression in CeA^PDYN+SST+^ neurons. Growth deficits were first detectable 5-7 days after the viral injection, further increase until ∼P17, and subsequently stabilize (**Fig 4g**, cyan and magenta traces). To determine whether growth deficits persist into adulthood, a third cohort of Pdyn-IRES-cre(+/-);Sst-IRES-flpO(+/-);Ai65(+/-) pups were injected with scAAV2/8-EF1a(core)-DDIO -DTA or -GFP into CeA at P1, and weights were monitored bi-weekly for 12 weeks. In this cohort, pups with DTA lesions of CeA^PDYN+SST+^ neurons recovered to ∼95% body weight of their -GFP injected littermates after weaning and maintained these relative body weights into adulthood (**Fig 4h**). Together these results demonstrate that the CeA^PDYN+SST+^ neurons are required for newborns to grow normally and thrive during the postnatal phase when pups are nursing, and that the weight deficit resulting from lack of this cell population largely recovers as independent feeding develops (*26, 72*).

Intriguingly, past work has shown that intracisternal (hindbrain) injections of dynorphin A in newborn rat pups increases attachment and suckling performance on an artificial nipple providing water (*22*). In addition, pairing dynorphin A with presentation of the nipple conditions subsequent attachment, consistent with a potential endogenous role of dynorphin in suckling behavior. Given these observations on the sufficiency of intracisternal Dynorphin A injections to enhance suckling, combined with our observations that the major output pathway of the CeA^PDYN+SST+^ neurons is to the hindbrain (**Fig 3),** we asked whether the ability of CeA^PDYN+SST+^ neurons to produce dynorphin is required for normal growth. We crossed Pdyn-floxed (Pdyn[fl](+/+)) mice (*73*) to Sst-IRES-cre(+/-);Pdyn[fl](+/-) mice, resulting in litters containing genotypes that include Sst-IRES-cre(+/-);Pdyn[fl](+/+) and Sst-IRES-cre(-/-);Pdyn[fl](+/+) littermates. We confirmed that *Pdyn* mRNA was eliminated in *Sst*-expressing neurons in the Sst-IRES-cre(+/-);Pdyn[fl](+/+) mice at P15 (**Fig 4i**). We did not observe significantly different body weights between these groups between p1 and p15 (**Fig 4j**), suggesting that *Pdyn* expression in CeA^PDYN+SST+^ neurons is not required for normal growth.

### CeA^PDYN+SST+^ neurons promote suckling vigor

Based on our data showing that CeA^PDYN+SST+^ neurons are active during suckling (**Fig 2**) and required for normal growth (**Fig 4**), we asked whether and how these neurons may modulate suckling behavior. Our data also demonstrated that specific aspects of suckling behavior, such as the sucking frequency and the propensity to attach to the dam (**Fig 1**) are malleable and state dependent, thus representing potential ways in which suckling may be modulated by forebrain influences. We performed a battery of behavioral assays to determine how suckling and other pup behaviors may be affected by ablating CeA^PDYN+SST+^ neurons (**Fig 5a**). To assess the pups’ propensity to both approach and attach to the dam at P 11-12, a timepoint in which CeA^PDYN+SST+^ neuron ablations already produced growth deficits (**Fig 4g**), we employed an assay in which pups are placed 5 cm from their anesthetized mother, prompting them to approach her ventrum and then attach to the nipple (**Fig 5a-c**). Pups with CeA^PDYN+SST+^ neuron ablations as well as littermate controls both appear to learn to approach the mother, as evidenced by the decrease in approach latency across subsequent trials within a behavioral session (Hazard ratio increases by a factor of 1.3 [95% c.i. 1.19-1.42] per trial for non-ablated pups and 1.16 [95% c.i. 1.06-1.26] for DTA-ablated pups) (**Fig S5**). However, pups with DTA-ablation of CeA^PDYN+SST+^ neurons displayed a significantly increased latency to attach (Hazard ratio 0.60 [95% c.i. 0.42-0.88], p = 0.008) but not approach the dam (Hazard ratio 0.92 [95% c.i. 0.62 1.36], p=0.68) when compared with non-ablated littermates (**Fig 5b,c**). Assessments of reflexive rhythmic sucking frequency using the artificial nipple assay (**Fig 1e**) at P9 showed that pups with DTA-ablations of CeA^PDYN+SST+^ neurons have a significantly reduced mean sucking frequency (frequency difference 8%, p=0.0014) but not amplitude (**Fig 5d**), as observed in sated versus deprived pups (**Fig 1h**). Together these results suggest that loss of CeA^PDYN+SST+^ neurons leads to specific decreases in suckling “vigor” (defined by frequency and inverse latency (*32, 33*)). Additionally, tests of the righting reflex at P9, a commonly used assay for mouse pup motor capability and strength, revealed that DTA-ablations of CeA^PDYN+SST+^ neurons resulted in longer latencies to right (Hazard ratio 0.62 [95% c.i. 0.41-0.95], p=0.028) (**Fig 5e**), which may be associated with a loss of strength associated with reductions in body weight and failure to thrive.

**Figure 5.**
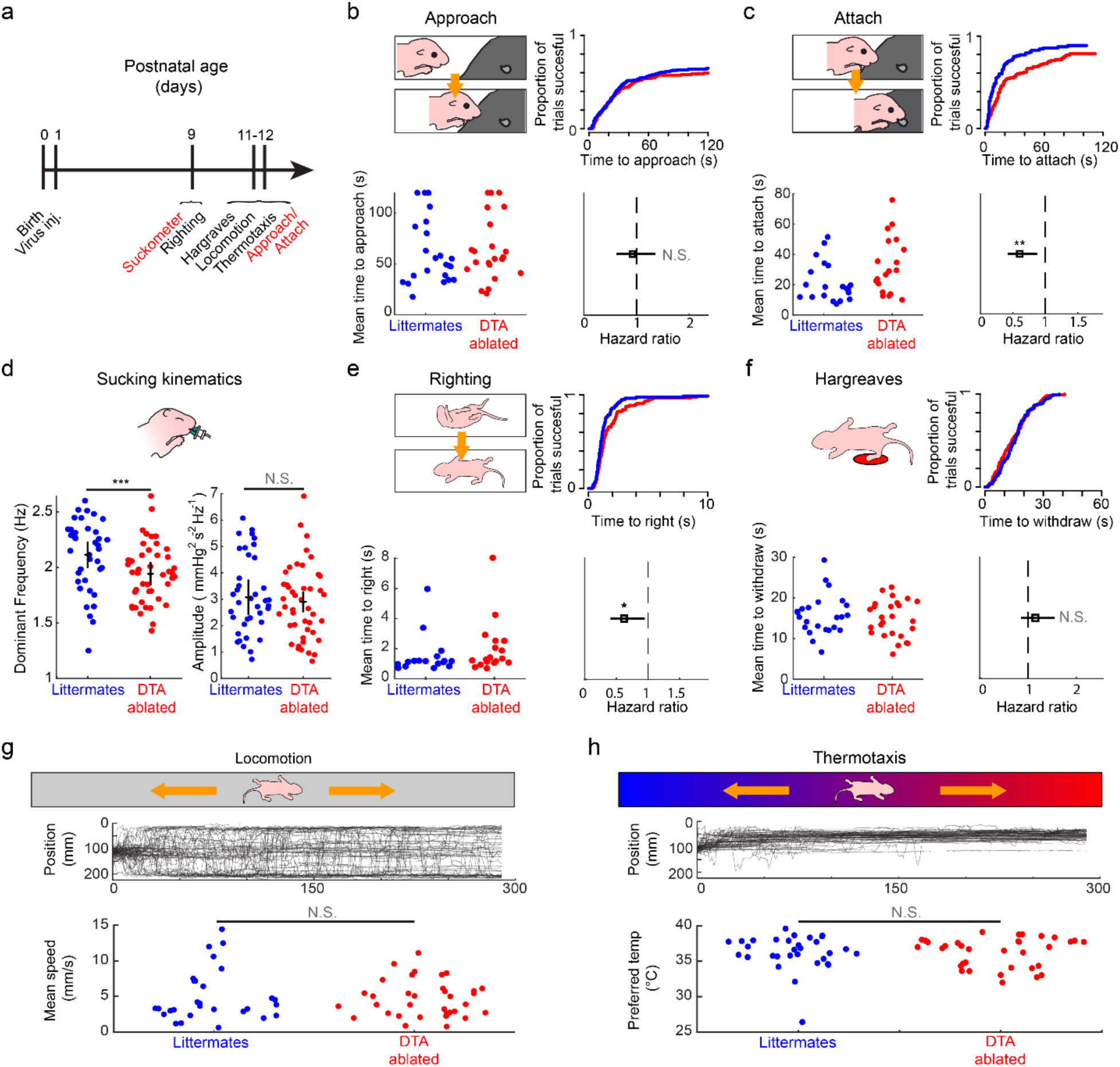
Rapid viral ablation of CeA^PDYN+SST+^ neurons leads to deficits in suckling vigor with minimal effects on other sensorimotor behaviors. **(a)** Timeline of assays for suckling-related and other behaviors for DTA-ablated versus non-ablated mouse pups (see Methods). These assays were performed on subsets of the litters represented in Fig 4g**. (b)** Approach and attachment assay: (top left) diagram of experimental measurement for approaching the mother: latency to reach the anesthetized mother’s ventrum when placed in a cage with her. (bottom left) Mean latency to approach over all trials per pup for DTA-ablated pups (red dots) and non-ablated littermates (blue dots). Pups that failed to approach within 120 s were assigned a latency of 120 s. (top right) Cumulative incidence curves showing the cumulative fraction of trials in which pups have approached at each latency. (bottom right) Hazard ratio (open square) and 95% confidence interval (line) for the propensity to approach for DTA-ablated versus non-ablated littermates. (N.S. - not significant, multilevel mixed-effects Cox regression accounting for individual and litter effects N=22 ablated pups and 23 non-ablated pups in 6 litters; see Methods). **(c)** Approach and attachment assay: Measurements of the latency to attach to the anesthetized mother’s nipple following successful approaches in **panel b**. Conventions are as in **panel b** (**p<0.01, multilevel mixed-effects Cox regression accounting for individual and litter effects) **(d)** (left) Mean dominant frequency of rhythmic sucking over all trials for each pup (***p<0.005 linear multi-level mixed-effects regression; N=45 ablated pups and 38 non-ablated pups from 12 litters). (right) Peak amplitude of rhythmic sucking for the same data (N.S. – not significant**). (e)** Righting reflex assay: measurements of the latency to right from supine to prone position. Pups that failed to right within 20s were assigned a latency of 20 s. Conventions are as in **panel b**. (*p<0.05, multilevel mixed-effects Cox regression accounting for individual and litter effects N=18 ablated pups and 17 non-ablated pups in 5 litters). **(f)** Nociceptive withdrawal assay: measurements of the latency to withdraw the paw from a thermal stimulus (Hargreaves assay). Pups that failed to right within 60s were assigned a latency of 60 s. Conventions are as in **panel b**. (N.S. not significant, multilevel mixed-effects Cox regression accounting for individual and litter effects, N=26 ablated pups and 24 non-ablated pups in 7 litters) **(g)** Locomotion assay: (top) Diagram of experimental measurement: speed of locomotion on a linear track at room temperature. (middle) Lines represent position versus time traces for each pup. (bottom) Mean speed of each DTA-ablated (red) and non-ablated (blue) pup. (N.S. – not significant, linear mixed-effects model for pups within litters; N= 34 ablated pups and 29 non-ablated pups in 9 litters) **(h)** Thermotaxis assay: (top) Diagram of experimental measurement: pup settling position on a linear track with a thermal gradient (middle). Lines represent position versus time traces for each pup. (bottom) Preferred temperature of each DTA-ablated (red) and non-ablated (blue) pup. (N.S. – not significant, linear mixed-effects model for pups within litters). This assay was performed on the same litters in **panel g** (N= 34 ablated pups and 31 non-ablated pups in 9 litters; see Methods).

To describe the broader range of behavioral changes that may be associated with the loss of CeA^PDYN+SST+^ neurons, we included other developmentally relevant assays of mouse pup behavior. These assays included responsiveness to a noxious heat stimulus to assess pain sensitivity and general motor capability (*74*), as well as locomotor and thermotaxis assays that could influence maternal seeking (*75*) (**Fig 5f-h**). We did not observe significant differences between DTA-ablated pups and control littermates in any of these assays (withdrawal latency to noxious heat: Hazard ratio 1.15 [95% c.i. 0.86-1.55], p=0.35; locomotor speed: 4.7 and 5.0 mm/s, respectively; p=0.68, linear mixed effects regression; preferred temperature: 36.27 and 36.28 deg C, respectively, p=0.98, linear mixed effects regression). Together, these results highlight that the behavioral deficits that result from ablation of the CeA^PDYN+SST+^ neurons are specific to modulating suckling, as opposed to more general behavioral deficits. The specificity of these deficits supports a specific role of CeA^PDYN+SST+^ neurons in maintaining the vigor of suckling throughout postnatal development.

## DISCUSSION

This study provides new insight into how the mammalian central nervous system regulates suckling behavior. We show that newborn mouse pups modulate nipple attachment and sucking rate, the defining characteristics of behavioral “vigor” (*32, 33*), according to their internal state (**Fig 1**). These data differ from previous results in rats in which nipple attachment was shown not to depend on milk deprivation until the second postnatal week (*11*),(*12*). This discrepancy may stem from inherent developmental or metabolic differences between rodent species; nonetheless, our observations demonstrate that newborn mice are capable of making value judgments and selecting actions accordingly essentially from birth. Next, we demonstrate that CeA^PDYN+SST+^ neurons show large increases in FOS expression when pups suckle (**Fig 2**). These neurons broadcast their axonal projections to diverse brainstem sites implicated in oral sensorimotor signaling and reinforcement learning, and they receive inputs from feeding-active hubs in the midbrain and hypothalamus (*76*) (**Fig 3**). Using new scAAV viral vector tools that enable cell-type-specific neuronal ablations in newborn mice, we then show that CeA^PDYN+SST+^ neurons are required for normal growth in early life (**Fig 4**), and that the loss of these neurons results in less vigorous nipple attachment and a reduced rate of rhythmic sucking (**Fig 5**).

Furthermore, this study provides evidence that infant suckling and adult feeding utilize separable neuronal circuit mechanisms, as hypothesized previously based on evolutionary and behavioral studies (*26*). Specifically, we observe that different populations of CeA neurons are active during newborn suckling versus adult feeding (**Fig 2**), and that CeA^PDYN+SST+^ neurons are required for body weight maintenance in early life but not in adulthood (**Fig 4**). Consistent with this hypothesis are the observations that (1) systemic ablations of *AgRP+* neurons, located in the arcuate nucleus of the hypothalamus, cause rapid starvation in adults but not neonates (*24*); (2) FOS activation in *AgRP+* neurons of P10 mice occurs after several hours of separation from the dam but is unrelated to nutrient deprivation (*77*); and (3) while chemogenetic activation of *AgRP*+ neurons influences pups’ approach to the dam at P10, it does not modulate nipple attachment or milk intake until P15 (*77*), when the transition to independent feeding begins (*78*). We note that a separate study indicates that more extreme levels of nutrient deprivation (16 hours) can lead to FOS activation in *AgRP*+ neurons at P10, and stimulation of these neurons optogenetically can lead to increased nipple attachment (*79*), suggesting that under certain physiological circumstances *AgRP*+ neurons could be recruited to promote suckling or maternal seeking behavior; however, they do not appear to be necessary when pups are reared in normal laboratory conditions (*24*). Together with our study, these observations strongly support the hypothesis that mammals possess separate but complementary feeding systems that are developmentally timed so that reliance on independent feeding ramps up as reliance on suckling winds down.

These previous studies on the roles of *AgRP+* neurons in early life also suggest that ethologically variable demands on infants may require neural substrates to adapt maternally-directed pup behaviors to different contexts and life circumstances. To sustain life, suckling must be executed reliably from birth, but must also be flexible enough to allow animals to survive in a variety of different environmental circumstances, such as adapting to differing maternal odors, varying milk availability, competition among littermates, and, in some contexts, bottle feeding (*9, 15, 17*). In addition to extracting milk, suckling triggers the dam’s milk release and production, and thus modulations in suckling vigor caused by CeA^PDYN+SST+^ neuronal activation may serve the immediate goal of acutely modulating milk intake, as well as the longer-term goal of altering the dam’s milk yield according to the pup’s needs (*80, 81*). Intriguingly, the amygdala is a highly conserved limbic brain structure that has long been shown to have a prominent role in implicit “emotional” learning, memory, and motivation (*28, 29, 82*). Indeed, recent studies have shown that: (1) stimulation of particular cell types, including *Sst*+ neurons, in the CeA can serve as a reinforcement signal for instrumental conditioning (*29*); (2) cell-type-independent stimulation of the CeA can intensify and narrow attraction to a particular reward target (*83*); and (3) SN-projecting CeA neurons, the majority of which express *Pdyn* and/or *Sst*, support conditioned reward-seeking (*40, 61*). Our findings are therefore generally consistent with previously demonstrated roles of the CeA in motivating behavior and supporting reward learning (*40*), and expand the known functions of the CeA to include supporting adaptations that occur during early life suckling.

Studies in a variety of species show that motor vigor is often associated with reward expectation (*32*). For example, primates trained to make saccades to a target location exhibit a lower latency to initiate movement and a higher velocity of movement when they expect a reward for saccading to the target (*84*). In this respect, motor vigor may serve as an overt readout of an animal’s subjective valuation of executing a movement or behavior (*32*). The observed decrease in suckling vigor resulting from either satiety (**Fig 1**) and/or the loss of CeA^PDYN+SST+^ neuron activity (**Fig 5**) may therefore reflect a decrease in the pups’ expected utility from suckling. If so, the activity of CeA^PDYN+SST+^ neurons during suckling might be itself reinforcing, consistent with observations that adult mice will work to receive stimulation of CeA^SST+^ neuron stimulation in an operant conditioning paradigm (*29*). The lack of such continued reinforcement to maintain the pups’ drive to suckle throughout infancy could account for the gradual body weight loss associated with CeA^PDYN+SST+^ neuron loss.

Finally, how might CeA^PDYN+SST+^ neurons exert their effects on downstream brain regions to influence suckling vigor? We note that the broad output projections from the CeA^PDYN+SST+^ include many brainstem premotor regions, which contain neurons that directly innervate oro-motor neurons (**Fig 3o**). Further, CeA^PDYN+SST+^ neuron projections to sensory input structures, e.g. the PB and NST, additionally may alter vigor by strengthening associations between specific sensory cues and expected reward, resulting in increased motivation to suckle. Altogether, these projections are poised to enhance different aspects of the suckling repertoire simultaneously across their diverse oro-motor, oro-sensory, and reward center targets.

## ACKNOWLEDGMENTS

We thank Prof. Martyn Goulding for the Rosa-DS-HTB mouse line, Huy Thong Phan for technical assistance with in-situ hybridization experiments, and Dr. Marito Hayashi for helpful discussions. Diagrams of experimental procedures were created using content from BioRender.com.

This work was supported by a Jane Coffins Child Postdoctoral Fellowship to J.D.M., NIH/NICHD grant K99/R00HD096512 to J.D.M, Kavli Institute for Brain and Mind Award KSIP-2017-001 to C.D., NIH/NICHD grant R01HD082131 to CD, and by the GT3 Core Facility of the Salk Institute with funding from NIH-NCI CCSG: P30 CA01495. C.D. is an investigator at the Howard Hughes Medical Institute.

J.D.M. and C.D. designed the experiments and wrote the paper. J.D.M. performed the experiments and analyzed the data. J.D.M., L.C.B., and S.L.P. designed and generated the viral vector constructs. J.D.M. and L.E.M. annotated neuroanatomical structures in histological sections. S.L.P. and C.D. provided research infrastructure.

## LIST OF SUPPLEMENTAL MATERIALS

Materials and Methods Figures S1-S5

