## supplemental material for "A genetically-defined population of amygdalofugal neurons promotes suckling and early postnatal growth"

### MATERIALS AND METHODS

#### Behavior tests

Attachment assay (P1)

Two groups of pups were tested for attachment to their anesthetized mother. The first group was placed in a clean cage on a heating pad set to 32deg C. The second group was placed in a clean cage with their mother. Both cages were supplied with mouse chow and hydrogel.

After 6 hours the mother was anesthetized with ketamine/xylazine and placed laterally recumbent on a heating pad. The pup was placed resting on a raised platform such that the head comfortably reached the nipple. The experimenter gently restrained each pup by the shoulders with their nose resting on the dam's nipple for 60 seconds. The pup/dam interaction was recorded on video (Point Grey Imaging Firefly MV USB camera). Videos were scored manually for the time at which the pup grasped the nipple with its mouth following placement on the dam's ventrum (*17*). For each trial, every pup in the litter was tested sequentially. The full assay consisted of 4-5 trials per pup.

During the assay the dam was monitored for responses to toe pinches, and given a supplemental dose of ketamine at 1/3 the initial dose when withdrawal responses to pinches were observed.

Oral gavage (P1)

A custom gavage device was constructed by attaching a thin flexible tube (24 gauge, 0.5mm OD) to a blunted hypodermic needle attached to a 1mL syringe (*85*) . Pups were isolated from their mother for 6 hours in a clean cage on a 32 deg C heating pad.

At the beginning of the assay, the gavage tube was inserted through the mouth approximately 1cm. In the sham group, the syringe was empty and no pressure was exerted on the plunger. In the milk group, approximately 0.05 mL of goat's milk esbilac (PetAg Inc.) were inserted by gently pushing on the plunger. Successful delivery of milk was confirmed by visualizing a milk spot in the stomach and weighing the pup before and after the procedure. Pups in which no milk was observed in the stomach or which the tube could not be inserted were immediately euthanized and not assayed.

For pups who successfully underwent the gavage procedure, we initiated the attachment assay described above.

Approach and attachment to anesthetized dam assay (P11-12)

The mother of a litter of P11-P12 pups was anesthetized with ketamine/xylazine and placed laterally recumbent at one side of a clean cage with standard bedding on a heating pad set to 32 deg C. Each test session consisted of each pup in a litter being tested for 8 trials lasting two minutes (120 s) each. Each pup in the litter was tested sequentially before moving to the next trial. In each trial, the pup was placed with their nose facing the dam, 5cm away from her ventrum. Pups' approach and attachment to the mother was video recorded from above (Logitech C270 webcam). The time to reach the mother, defined as the nose occluding the mother's ventrum in the video, as well as the time to grasp the mother's nipple were recorded. The latency to approach was defined as the time between the start of the trial and the pup reaching the mother's ventrum. The latency to attach was defined as the time between a successful approach and nipple grasping.

Sucking pressure assay (P3-P9)

Pups aged P3-P9 were removed from their home cage and placed on a heating pad immediately prior to the assay. Before each trial, the empty silicone nipple was dipped in sunflower oil to provide a mechanical sealant (*86*) and blotted on a paper towel. In each trial, the experimenter held the pup gently by the nape and inserted the empty silicone nipple in the mouth, between the tongue and the palate to a depth such that the nose was touching the plastic stopper (**Fig 1**). Pups reliably initiated sucking movements. The nipple was held in the mouth for 60 seconds. In trials where the pup rejected the nipple it was immediately re-inserted. The nipple, connected to a pressure transducer and amplifier (see **Mouse Pacifier Fabrication**), was used to monitor intraoral pressure during rhythmic non-nutritive suckling. Pressure waveforms were digitized using a Powerlab analog-to-digital converter (AD Instruments) at 100 samples/second. For each trial, every pup in the litter was tested sequentially. The full assay consisted of 4-5 trials per pup.

Locomotion/thermotaxis assay (P9-P10)

*Locomotion:* Each test session consisted of each pup in a litter being tested for one trial lasting five minutes each. In each trial, the pup was placed in the center of a walled, linear aluminum track at room temperature (23cm x 4cm x 0.2cm). Pups' locomotion on the track was monitored on video (Microsoft Lifecam webcam).

*Thermotaxis:* Following the locomotion assay, pups were again tested on the linear track for five minutes each. For this assay, a thermal gradient was applied to the linear track with one end being held at 10 degrees C using a custom made, water-recirculating Peltier cooler (GeeBat TEC1-12706 Thermoelectric Cooler 12V 5.8A, Amazon Inc; BXQINLENX Aluminum Water Cooling Block for CPU Graphics Radiator Heatsink 40x 40mm, Amazon Inc.) and thermostat (Baylite BTC201 Pre-Wired Digital Outlet Thermostat, Amazon Inc.), and the other end held at 40 degrees C using a custom made nichrom wire heating coil and thermostat. Temperature along the track was measured with thermistors (LM-35, Digikey Inc) placed at 5 equally spaced points along the track and sampled via a microcontroller (Arduino Uno). Pups' locomotion on the track was monitored on video (Microsoft Lifecam webcam).

For both locomotion and thermotaxis assays, video and real-time temperature information were recorded simultaneously via custom scripts written for the Bonsai programming platform (*87*) . Video images were thresholded to detect the center of mass of the pup in each frame. The locations of the temperature sensors were annotated manually, and the temperature corresponding to the position of the pup was estimated by linear interpolation between the positions of the temperature sensor readings. To calculate locomotor speed, we used the magnitude of the velocity of the pup's center of mass averaged over the full five-minute trial. To calculate the preferred temperature, we used the average temperature corresponding to the pup's center of mass over the final 60 seconds of the assay, at which point pups tended to remain stationary when the temperature gradient was on (**Fig 5h**). Video recordings failed due to software freezing in 3 of 130 trials (2%).

<u>Righting reflex assay (P9-P10)</u>

Each test session consisted of each pup in a litter being tested for 5 trials lasting a maximum of 20 seconds each. Each pup in the litter was tested sequentially before moving to the next trial. In each trial, the pup was held supine on a plastic weighing dish, pinned by the experimenter's thumb and index fingers holding the pup's forelimbs and hindlimbs. The pup was then released, and the time to right, defined as the flipping from the supine to the prone position with all paws touching the ground (*88*), was recorded.

Nociceptive withdrawal (Hargreaves) assay (P9-P10)

Each test session consisted of each pup in a litter being tested for 5 trials lasting a maximum of 60 seconds each. Each pup in the litter was tested sequentially before moving to the next trial. In each trial, pups were placed in the prone position atop a Thermal Plantar Test apparatus (Stoelting # 55370) inside a clear acrylic chamber. The supplied infrared heat source (30 mW/cm^2) was placed underneath the glass floor of the apparatus, directly under the hindpaw of the pup, and the time to withdraw the paw from the floor of the test apparatus was automatically detected and recorded.

Immediate early gene assays

Pups were put through the following behavioral protocols to assess FOS activity:

*Suckling/Isolation:* On each test day, 4-6 P2-P3 pups from a single litter were weighed, isolated from their mother for 6 hours, reweighed, and then placed in a clean cage on a heating pad set to 32 deg C. Their mother was placed alone in a separate clean cage and given food and hydrogel. After 6 hours, 2-3 of the isolated pups were placed in the mother's cage and the interaction was monitored with video recording (Logitech C270 webcam). We observed for continuous nursing bouts of at least 20 minutes, which typically occurred within 0-2 hours of the reunion. For immunohistochemical assays, pups were weighed, cryoanesthetized, and decapitated 80 minutes following the onset of the nursing bout, and the heads were immediately placed in iced 4% paraformaldehyde. For in-situ hybridization assays, pups were weighed, cryoaneshetized, and decapitated 30-35 min following the onset of the nursing bout, and the heads were frozen in OCT on dry ice. In all cases, the remaining isolated, non-nursed littermates were sacrificed at the same time using the same procedure.

*Huddling/Isolation:* On each test day, 4-6 P2-P3 pups from a single litter were weighed, isolated from their mother for 6 hours, reweighed, and then placed in a clean cage on a heating pad set to 32 deg C. A non-lactating female who had previous maternal experience was placed alone in a separate clean cage and given food and hydrogel. After 6 hours, 2-3 of the isolated pups were placed in the non-lactating female's cage and the interaction was monitored with video recording (Logitech C270 webcam). We watched for continuous maternal bouts, where the female huddled over and licked the pups continuously for at least 20 minutes, which typically occurred within 0-2 hours of the reunion. In no case did we observe the pups attach to the female's nipple. Pups were subsequently weighed, cryoanesthetized, and decapitated 80 minutes following the onset of the huddling bout, and the heads were immediately placed in iced 4% paraformaldehyde.

*Agitation:* On each test day, P2 pups from a single litter were weighed, isolated from their mother for 6 hours, reweighed, and then placed in a clean cage on a heating pad set to 32 deg C. After 6 hours, the experimenter manually handled and massaged the pups in their cage for 60 minutes and the procedure was recorded on video (Logitech C270 webcam). For immunohistochemical assays, pups were cryoanesthetized and decapitated 80 minutes following the onset of the handling procedure, and the heads were immediately placed in iced 4% paraformaldehyde.

*Gavage feeding:* On each test day, P2-P3 pups from a single litter were weighed, isolated from their mother for 6 hours, reweighed, and then placed in a clean cage on a heating pad set to 32 deg C. After 6 hours, the experimenter inserted a feeding tube and delivered either 100uL of GME or performed a sham insertion in which no liquid was delivered, according to the procedure described above (see Oral Gavage). For immunohistochemical assays, pups were cryoanesthetized, and decapitated 80 minutes following the gavage procedure, and the heads were immediately placed in iced 4% paraformaldehyde.

*Intraoral milk delivery:* On each test day, P2-P3 pups from a single litter were weighed, isolated from their mother for 6 hours, reweighed, and then placed in a clean cage on a heating pad set to 32 deg C. After 6 hours, the experimenter inserted a feeding tube (see Oral gavage) ~1-2 mm into the mouth between the tongue and the palate and dispensed small volumes of Goat's milk Esbilac (PetAg) into the mouth until liquid was observed to spill out of the mouth. Excess liquid was blotted with a paper towel. The procedure was repeated at a duty cycle of 3-5 minutes every 10 minutes for 80 minutes. For immunohistochemical assays, pups were cryoanesthetized, and decapitated 80 minutes following the first intraoral delivery, and the heads were immediately placed in iced 4% paraformaldehyde.

<u>FosTRAP assays:</u>

Homozygous Fos-creER (TRAP2) mice (*41*) were bred with homozygous LSL tdTomato (tdT) mice (Ai9 or Ai14) to produce TRAP +/-; LSL tdT +/- pups . On P2 or P3, pups were injected with 50mg/kg 4-hydroxy-tamoxifen (prepared as 25mg/mL in 100% EtOH and diluted 1:10 in sunflower oil). In some experiments, pups were put through the suckling immediate early gene assay described above on P6, and the brains were prepared for immunohistochemical processing. In other experiments, pups were allowed to grow to adulthood (2-6 months, and subsequently underwent one of the following immediate early gene assays:

*Feeding:* Prior to the assay mice were weighed, single housed and food deprived for 23 hours, and weighed again. At the start of the trial mice were given chow coated with a palatable high-calorie supplement (Nutri-cal, Vetoquinol USA) and monitored via video recording. All mice were observed to immediately begin to consume the food. Littermate control mice remained food deprived throughout the assay. After 80 minutes, mice were weighed again, anesthetized with carbon dioxide exposure and underwent transcardial perfusion.

*Sucrose:* Prior to the assay mice were weighed, single housed, water-restricted for 23 hours, and weighed again. At the start of the trial mice were given water bottles containing a 2% sucrose solution with a known weight. All mice were observed to immediately begin to consume the sucrose solution. Littermate control mice remained water-deprived throughout the assay. After 80 minutes, mice and water bottles were weighed again, and mice were anesthetized with carbon dioxide exposure and underwent transcardial perfusion.

**Viral vector creation:**

*scAAV-Flex-GFP:* To rapidly trace the anatomical projections originating from specific cell types, we used a previously reported self-complementing scAAV-Flex-GFP plasmid (*57*) . To generate virus, the plasmid was transformed into chemically competent e.coli cells (OneShot Stbl 3, Invitrogen) according to manufacturer specifications and amplified with a maxiprep kit (Invitrogen) according to manufacturer specifications. Plasmid DNA was provided to the Salk Institute GT3 core to generate AAV2/8 vectors with titers ranging from 10^13 to 10^15 genome counts/mL.

*scAAV-FLEx-synaptophysin-HA:* We modified the scAAV-FLEx-GFP plasmid to express HA-tagged synaptophysin instead of GFP, using standard molecular cloning techniques. Briefly, we first created an open scAAV-Flex backbone by using restriction enzymes NcOI-HF and BsrGI (New England Biolabs). We then designed primers to excise the human synaptophysin gene from an existing AAV plasmid (*89*), and included the HA sequence in the forward primer. The synaptophysin-HA construct was then excised and replicated using the Phusion PCR system (New England Biolabs) according to manufacturer specifications.

The digested scAAV backbone and synaptophysin-HA PCR products were then run on an electrophoresis gel, and the bands corresponding to these constructs were extracted using a gel extraction kit (Macherey-Nagel Inc.) according to manufacturer specifications. The synaptophysin-HA DNA was cloned into the scAAV backbone using the In-Fusion Cloning Kit (Takara Biosci.) according to manufacturer specifications.

*scAAV-DDIO-GFP and scAAV-DDIO-DTA:*

We synthesized DNA constructs, “gBlocks” (Integrated DNA Technologies Inc.), containing unidirectional *lox* recombination site mutants for cre-dependent expression (*lox71* and *loxJTZ17*), unidirectional *frt* recombination site mutants for flp-dependent expression (*frt-LE(-10)* and *frt-RE(+10)*), and modified splice donor/acceptor sites, as well as the DTA or GFP fragment segments to reconstitute DTA and GFP proteins, respectively, in a cre- and flp- dependent manner. The segments were cloned into an scAAV-EF1a backbone as described above. The fragment segments were designed such that the expression of one fragment alone in the presence of either cre- or flp-, but not both, would not yield a functional DTA or fluorescent GFP.

**Virus injections:**

<u>AAV injections:</u>

For cell-type-specific anterograde anatomical tracing and ablations we injected self-complementing adeno-associated viral (scAAV) vectors into the central amygdala on postnatal day 1 (P1). For retrograde tracing, we injected dG-rabies virus in the same location. Briefly, mouse pups were cryoanesthetized and placed in a stereotaxic holding frame atop a bed of ice. A layer of silly putty was affixed to the ends of zygomatic ear bars, which were used to gently hold the head in the stereotaxic frame. The blood sinus under the lambda suture was visualized through the skin and used as a stereotaxic reference point. The virus was pressure injected at 1.4 mm rostral, 1.7mm lateral, to the lambda suture, and 3.2 mm ventral to the brain surface. For each injection, virus was pressure-expelled through a glass micropipette (WPI BF100F4) with a 20-30um tip.

*Anterograde anatomical tracing:* A mixture containing cre-dependent scAAV2/8-hSyn-FLEx-GFP at 1 x 10^13 GC/mL and cre-dependent scAAV2/8-hSyn-FLEx-Syp-HA at 1 x 10^13 GC/mL (Salk GT3 Core) was pressure injected (15ms pulses, 5-20 PSI, approximately 30-40 nL total volume) unilaterally into the brain of mouse pups with genotype Pdyn-ires-cre(+/-) (Jax stock no. 027958):Sst-ires-flpO (+/-) (Jax stock no 028579) x RCFL-tdT (+/-) (Jax stock no 021875). In some experiments, the cortical area dorsal to the injection site was removed to limit viral uptake by sparsely-labeled Pdyn+, Sst+ cortical neurons along the pipette track. After 7 days, the mice were deeply anesthetized with CO2 and decapitated.

*Retrograde anatomical tracing:* EnvA-pseudotyped SADB19-dG-Rabies-mCherry virus at 10^9 GC/mL (Salk GT3 Core) was pressure injected (15ms pulses, 5-20 PSI, approximately 120 nL total volume) unilaterally into the brain of mouse pups resulting from crosses of a Pdyn-ires-cre (+/+) Sst-iresFlpO (+/+) parent and a cre- and flp-dependent (Los-Stop-Lox-Frt-Stop-Frt) histone tagged-GFP (H2B-GFP), TVA, and Rabies-glycoprotein (B19G) expressing parent mouse (Rosa-DS-HTB); +/+; courtesy Martyn Goulding, Salk Institute) (*63, 64*). After 7 days, the mice were deeply anesthetized with CO2 and decapitated.

*Cell-type-specific ablation:* Pups were injected bilaterally with cre- and flp-dependent scAAV2/8-EF1a(core)-DDIO-DTA at 1 x 10^14 GC/mL (Salk GT3 Core, 15ms pulses, 5-20 PSI, approximately 60 nL per side). In some experiments, litters resulting from crosses of a Pdyn-ires-cre(+/-);Sst-iresFlpO(+/+) or Pdyn-ires-cre(+/+);Sst-ires-FlpO (+/-) parent and a RCFL-tdT parent (+/+) mouse were used, and the entire litter was injected on P1. These litters therefore consisted, on average, of half the pups in each litter with both the

Pdyn-ires-cre and the Sst-ires-FlpO allele. In other experiments, litters resulting from crosses of a Pdyn-ires-cre (+/+);Sst-iresFlpO (+/+) parent and a RCFL-tdT parent (+/+) mouse were used, and half the pups in the litter were injected with scAAV2/8-EF1a(core)-DDIO-DTA at 1 x 10^14 GC/mL and the other half of the pups were injected with scAAV2/8-EF1a(core)-DDIO-GFP at 1 x 10^14 GC/mL. Pups' body weights were monitored and a subset of the behavioral assays described above, including approach/attach, sucking pressure, locomotion/thermotaxis, and righting reflex were performed between P9 and P12 (see main text and **Behavior tests** section). In some experiments, mice were transcardially perfused with 4% PFA at the end of the monitoring period. Pups were occasionally found dead following virus injections, independent of virus type/genotype (6 ablated and 6 non-ablated out of 140 injected mice), and excluded from subsequent body weight measurements.

**Histology**

Immunohistochemistry:

*Tissue preparation:* For pups < P14, heads were fixed in 4% PFA for 2-3 days, after which the brains were dissected and postfixed for 1 additional day.

For pups >= p14, mice were transcardially perfused with 4%PFA using a manual syringe. Heads were post-fixed in 4% PFA for 2-3 days, after which the brains were dissected and postfixed for 1 additional day.

For all experiments, brains were sectioned into 50-60 um sections with a sliding microtome or cryostat and collected in phosphate buffered saline (PBS) (Corning or Gibco). Sections were washed one time in PBS, incubated for 20 minutes at room temperature in blocking solution (2% Normal Horse Serum (Vector Labs), 0.2% Triton X in PBS), and then incubated overnight in primary antibody solution (primary antibody in blocking solution). Sections were then washed 2 times in PBS and incubated secondary antibody solution (secondary antibody in blocking solution, washed another one time, and then mounted on glass slides.

One or more of the following primary antibodies were used:

*FOS:* For mouse pups, polyclonal rabbit anti-cFos (Cedarlane #226003(Sy), 1:2000) was used. For adult mice, either polyclonal rabbit anti-cFos (Cedarlane #226003(Sy), 1:2000) or monoclonal rabbit anti-cFos(9F6) (1:1000; Cell Signaling Technology #2250) was used. *anti-GFP:* Polyclonal chicken anti-GFP (Aves Labs GFP1020, Fisher Sci NC9510598, 1:1500) was used. *anti-RFP:* Polyclonal rabbit anti-RFP (MBL International, Fisher Sci #5050699, 1:1000) was used. *anti-HA*: Monoclonal mouse anti-HA.11 Epitope tag (Biolegend #901513, 1:1000) was used. *Anti-NeuN:* Polyclonal rabbit anti-NeuN (Millipore Sigma ABN78, Fisher Sci #ABN78MI, 1:1000) was used.

The following secondary antibodies was used at a concentration of 1:500: Alexa Fluor 647 donkey anti-rabbit IgG (Life Technologies #A31573); Alexa Fluor 488 donkey anti-rabbit IgG (Life Technologies #A21206); Alexa Fluor 594 donkey anti-rabbit IgG (Life Technologies A21207); Alexa Fluor 488 goat anti-chicken (Life Technologies #A11039); Alexa Fluor donkey anti-mouse IgG (Life Technologies #A32787).

In some experiments, sections were counterstained with Neurotrace Blue (N21479, 1:200) or Neurotrace Deep Red (Life Technologies N21483, 1:200) following manufacturer specified protocols.

In-situ hybridization:

*Tissue Preparation:* P2 mouse pups were decapitated and heads were immediately frozen in OCT on dry ice and stored at -80 degrees C. Immediately, prior to sectioning, the frozen tissue was equilibrated at -20 degrees C in the cryostat for >20 min. Brains were then sectioned into 20um sections on a cryostat and collected on glass slides.

*RNAscope and Hybridization Chain Reaction:* Sections were processed with the following RNAScope or Hybridization Chain Reaction (HCR) reagents according to manufacturer specified protocols: RNAscope Probe-Mm-Fos (Advanced Cell Diagnostics #316921), RNAscope Probe-Mm-Pdyn-C2 (Advanced Cell Diagnostics #318771-C2); RNAscope Probe-Mm-Sst-C3 (Advanced Cell Diagnostics #404631-C3); RNAscope Probe-Mm-Pnoc (Advanced Cell Diagnostics #437881); RNAscope Probe-Mm-Fos-O2-C2 (Advanced Cell Diagnostics #506921-C2); RNAscope Probe-Mm-Fos-O3-C2 (Advanced Cell Diagnostics #506931-C2); RNAscope Probe-Mm-slc32a1-C3 (Advanced Cell Diagnostics #319191-C3); RNAscope Multiplex Fluorescent Reagent kit (Advanced Cell Diagnostics #320850); HCR probes Slc32a1-B2, SST-B4, PDYN(exon 4)-B5 (custom) with amplifier fluorophores B2-488, B4-546, B5-647 (Molecular Instruments Inc).

*ISH with RNA probes:* For in-situ hybridization with RNA probes, we followed previously described techniques (*54*). Briefly, Complementary DNA of *Fos* and CeA molecular markers were cloned in approximately 800-base-pair (whenever possible) segments into pCRII TOPO vector (Invitrogen). Antisense cRNA probes were synthesized with T7 or Sp6 polymerases (Promega) and labeled with digoxigenin (DIG; Roche) or fluorescein (FITC; Roche). Tissue sections were processed as described previously and incubated with the RNA probes (*54*). Probes were subsequently revealed with peroxidase-conjugated antibodies (anti-DIG-POD or anti-FITC-POD, respectively, Roche) and fluorophore- conjugated peroxidase substrates (FITC-TSA, Cy3-TSA, or Cy5-TSA; Perkin Elmer).

**Data analysis:**

Microscopy:

*Slide scanning:* Microscopy slides were imaged using and Axioscan 7 slide scanner with a 10x/0.45NA objective for detecting somata (0.65um/pixel) and 20x/0.8NA objective (0.325um/pixel) for detecting boutons.

*Image analysis:* Slide scanning images were processed using custom written scripts in MATLAB (Mathworks, Natick, MA) or standard image segmentation algorithms in quPath (*90*) to identify somata and presynaptic terminals based on their size, shape, and fluorescent intensity. Neuroanatomical regional boundaries were annotated manually based on a reference atlas for neonatal mice (*91*) using fluorescent Nissl staining, anti-NeuN immunohistochemistry, and/or autofluorescence. Segmented cells or nuclei were defined as co-localized if their centroids fell within 10um of each other.

Statistical analysis:

*Calculation of sucking frequency and amplitude with the artificial nipple assay:* Power spectra were estimated over the full duration of each artificial nipple suckling trial using the multi-taper method (*92*) with a time-bandwidth product of 5 and 9 tapers, over the frequency range of 0.5 – 5 Hz. Spectra were calculated on the one-sample time derivative of the intraoral pressure signals. The Chronux package for MATLAB was used for spectral estimation (*93*) . The dominant sucking frequency in each trial was estimated by first subtracting the baseline power from the spectrum, defined as the mean power over all frequencies, and then calculating the center of mass of the portion of the mean-subtracted spectrum that exceeds 50% of the peak spectral power. The peak suckling amplitude is defined as the power at estimated dominant frequency.

*Linear multilevel mixed effects model:* For statistical analysis of behavioral tests we used multi-level mixed effects regression models to account for correlated observations both within individual subjects and within litters. Statistical calculations were performed using MATLAB or R software. Specifically, for behavioral observations measuring body weight, locomotor speed, and preferred temperature, the following model was used:

$$y \sim x + (1|LitterID)$$

where *y* represents the behavioral observation, *x* is a dummy variable representing the treatment or genotype, and *litterID* is an index variable that denotes which observations arise from animals from the same litters.

For behavioral observations that involved repeated measurements in each individual pup, including measurements dominant sucking frequency and amplitude, the following model was used in the MATLAB Statistics Toolbox:

$$y \sim x + (1|litterID) + \ (1|LitterID{:}\,mouseID)$$

where, *mouseID* is an index variable denoting which observations arise from the same individual animal.

For behavioral observations measuring latency to an event we used a mixed-effects Cox proportional hazards regression model in the R 'coxme' package (*94*):

$$surv(t, censor) \sim x + (1|\ litterID/mouseID\ )$$

where *surv(t)* represents the latency to the observed event, censored if the event was not observed within the observation period.

To assess whether latency to an event changes with learning across trials in a behavioral session we used a mixed-effects Cox proportional hazards regression model:

$$surv(t, censor) \sim trialNumber + (1|\ litterID/mouseID\ )$$

where *surv(t)* represents the latency to the observed event, censored if the event was not observed within the observation period, and *trialNumber* is the integer corresponding to the number of the trial within the behavioral session.

In all cases where the Cox proportional hazards model was utilized we first verified the validity of the proportional hazards assumption using the 'cox.zph' function in R.

*Generalized linear mixed effects model:* For statistical analysis of labeled cells we used binomial mixed effects regression models to estimate the proportions of double-labeled cells and confidence intervals around these proportions, accounting for correlated observations within subjects.

**Mice**

Mice were purchased or derived from Jackson Laboratory stocks as noted and maintained on a 12hr light cycle in standard laboratory vivarium conditions. All experiments were performed in accordance with IACUC regulations at Harvard University (Protocol 97-03), the Salk Institute (Protocol 11-00020), and the University of Southern California (Protocol 21505). Pup age estimates are accurate to +/- 1 day.

**Mouse pacifier fabrication:**

The mouse pacifier was constructed by filling the inside of a 0.2mL PCR tube with silicone elastomer (Body double TM fast-set). The bottom 5-6 mm was used for the nipple. A ~1mm hole was bored through the center of the silicone nipple, and a 1/16” plastic barbed leur fitting was threaded through the hole. This construction yielded a conical nipple that is 4-5 mm in diameter at the base and 5-6mm in length. The leur was connected via PTFE (Teflon) tubing to a pressure transducer (SMI #5652-001) and amplifier (AD627) with a gain of 25x.

### SUPPLEMENTAL FIGURE LEGENDS

**Figure S1. FOS antibody staining in various behavioral conditions and brain areas in suckling and non-suckling behavioral conditions.**

**(a)** FOS antibody staining (green) in the NST following suckling (left), huddling with a non-lactating dam (middle), or after gavage feeding with goat's milk esbilac (right) in a P2 pup. **(b)** FOS antibody staining (green) in the hypothalamus following suckling (left), huddling with a non-lactating dam (middle), or after being manually agitated by the experimenter (right) in a P2 pup. **(c)** (left) Mean density of FOS-expressing cells in suckling pups (blue bars) and huddling pups (gray bars) for regions with substantial anti-FOS immunolabeling throughout the brain. Measurements in individual pups are shown as black dots. (right) Difference in mean FOS density between suckling and huddling conditions. Anatomical abbreviations: Central amygdala (CeA), Nucleus of the solitary tract (NST), interstitial nucleus of the posterior limb of the anterior commissure (IPAC), centromedian thalamus (CM), lateral parabrachial area (LPB), piriform cortex (PIR), anterior nucleus of the paraventricular thalamus (PVA), striatum (Str), bed nuceus of the stria terminalis (ST), basolateral amygdala (BLA), median preoptic nucleus (MnPO), olfactory tubricle (Tu), nucleus accumbens (Acc), external cuneate (ECn), extended amygdala (EA), parvocellular reticular formation (PcRT), cuneate nucleus (CUN), medial amygdala (MeA), lateral periacqueductal gray (LPAG), septum (Sep), medial preoptic area (MPA), insular cortex (ins), ventral pallidum (vp), lateral hypothalamus (LH), globus pallidus (gp), posterior hypothalamus (PH), vascular organ of the lamina terminalis (VOLT), arcuate nucleus (arc), medial division of the central amygdala (CeM), lateral division of the central amygdala (CeL), capsular division of the central amygdala (CeC). **(d)** FOS immunolabeling (green) in the CeA following suckling (left) or isolation (right) in a P6 pup. **(e)** FOS immunolabeling (green) in the CeA following suckling (left) or isolation (right) in P11 pup. **(f)** Representative image of an adult TRAP2;Ai14 (TRAP) mouse CeA after injection of 4-OHT on P3 and anti-FOS immunolabeling after consuming 2% sucrose in adulthood. tdTomato expression is shown in red (left), FOS immunolabeling in green (middle), and a merge of these two images (right). All scale bars are 500um. Anatomical abbreviations: Nucleus of the solitary tract (NST), area postrema (AP), dorsal motor nucleus of the vagus (10N), lateral hypothalamus (LH), dorsomedial hypothalamus (DMH), ventromedial hypothalamus (VMH), arcuate nucleus (Arc), central amygdala (CeA) lateral (CeL), central amygdala medial (CeM), central amygdala capsular (CeC), basolateral amygdala (BLA), intercalated nucleus (I), striatum (Str).

**Figure S2. Evaluation of candidate molecular markers for suckling-active CeA neurons.**

**(a)** In-situ hybridization for *Fos* mRNA (red) and pro-dynorphin mRNA (green) in P2 a mouse pup following suckling (top) or isolation (bottom). **(b)** In-situ hybridization for *Fos* mRNA (red) and either *vGat* mRNA (green) (top left), *dlk1* mRNA (green) (bottom left), *nts* (neurotensin) mRNA (green) (top right), *pnoc* mRNA (green) bottom right in P2 mouse pups following suckling. **(c)** Images of RNAscope (in-situ hybridization) labeling of *Fos* (top), *Pdyn* (2$^{nd}$ from top), *Sst* (3$^{rd}$ from top) mRNA in the CeA following suckling at P2, and an overlay of these images (bottom, replication of **Fig 2I** for comparison) **(d)** Hybridization chain reaction (HCR®) in-situ hybridization for *Pdyn* mRNA (green), *Sst* mRNA (blue), and *vGat* mRNA (red) at P8. **(e)** Enlarged view of the boxed region in **panel**

**d** (top) with all fluorescent channels displayed, (middle) with only *Pdyn* and *Sst* channels, and (bottom) with only *Sst* and *vGat* channels **(f)** Image of a P2 Sst-ires-cre(+/-);Ai14(+/-) mouse CeA with anti-FOS immunolabeling after suckling. FOS immunolabeling is shown in green (left), tdTomato expression in red (middle), and a merge of these two images (right). **(g)** Image of a P2 Sst-ires-cre(+/-);Ai14(+/-) mouse CeA with anti-somatostatin immunolabeling. SST immunolabeling is shown in green and tdTomato expression in red. **(h)** Image of an adult Sst-ires-cre(+/-);Ai14(+/-) mouse CeA following an injection of scAAV2/8-hSyn-FLEx-GFP on P1. GFP expression is shown in green and tdTomato expression in red. **(i)** Image of a P2 Pdyn-ires-cre(+/-);Ai14(+/-) mouse forebrain at the level of CeA. tdTomato expression is shown in white. **(j)** Image of a P2 Sst-ires-cre(+/-);Ai14(+/-) mouse forebrain at the level of CeA. tdTomato expression is shown in white. **(k)** (top) Counts of $CeA^{PDYN+SST+}$ neurons (red) and FOS immunolabeled cells (green) in CeA subdomains of Pdyn-IRES-cre(+/-);Sst-IRES-flpO(+/-);Ai65(+/-) mice following suckling on P2. (bottom) Proportion of cells co-expressing FOS and tdTomato (N=4). All scale bars are 500um. Anatomical abbreviations: central amygdala (CeA) lateral (CeL), central amygdala medial (CeM), central amygdala capsular (CeC).

**Figure S3. 3D reconstructions of $CeA^{Pdyn+Sst+}$ input/output connectivity.**

**(a)** Whole brain reconstruction of terminals triple-labeled with anti-GFP, anti-tdTomato, and anti-HA (yellow) following the injection strategy shown in **Fig 3b**. The reconstruction is shown at an oblique view, replicated from **Fig 3i**, superimposed with outlines of the anatomical structures annotated in **Fig 3j** (red, green, blue, magenta, cyan) in serial sections (left), and a frontal view of the same reconstruction (right). **(b)** Sample whole brain reconstruction of CeA$^{PDYN+SST+}$ starter cells (green) and neurons identified as presynaptic to them (red) following the injection strategy in **Fig 3k**.

**Figure S4. Technical evaluation of the specificity of scAAV-DDIO vectors.**

**(a)** Images of a coronal section through the CeA of a P6 Pdyn-IRES-cre(+/-);Sst-IRES-flpO(+/-);Ai65(+/-) mouse with an injection of scAAV2/8-EF1a(core)-DDIO-GFP targeted to the CeA at P1. tdTomato expression is shown in red (left), GFP expression is shown in green (middle), and a merge of the two images (right). **(b)** Sample images of a coronal section through the CeA of a P6 Pdyn-IRES-cre(+/-);Sst-IRES-flpO(-/-);Ai65(+/-) mouse, a littermate of the mouse in **panel a** with an injection of scAAV2/8-EF1a(core)-DDIO-GFP targeted to the CeA at P1. (Lack of) tdTomato expression is shown in red (left), (lack of) GFP expression is shown in green (middle), and a merge of the two images (right). All scale bars are 500um. Anatomical abbreviations: central amygdala (CeA) lateral (CeL), central amygdala medial (CeM), central amygdala capsular (CeC). **(c)** (left) Body weights measured every two days from P1 to P28 following P1 injections of scAAV2/8-EF1a(core)-DDIO-GFP (blue, N=29 from 8 litters) or -DTA (red, N=27 littermates) targeted to the CeA in Pdyn-IRES-cre(+/-);Sst-IRES-flpO(+/-);Ai65(+/-) mouse pups. In each litter, approximately half the pups were injected with each vector. This is a replication of **Fig 4f** for reference. (middle) Body weights measured every two days from P1 to P28 following P1 injections of scAAV2/8-EF1a(core)-DDIO-DTA targeted to the CeA in Pdyn-IRES-cre(+/-);Sst-IRES-flpO(+/-);Ai65(+/-) mouse pups (red, N=12 from 4 litters) and Pdyn-IRES-cre(-/-);Sst-IRES-flpO(+/-);Ai65(+/-) littermates (blue, N=14). (right) Body weights following P1 injections of scAAV2/8-EF1a(core)-DDIO-DTA targeted to the CeA in Pdyn-IRES-cre(+/-);Sst-IRES-flpO(+/-);Ai65(+/-) mouse pups (red, N=19 from 4 litters) and Pdyn-IRES-cre(+/-);Sst-IRES-flpO(-/-);Ai65(+/-) littermates (blue, N=9). Summary statistics for these data are shown in **Fig 4g.**

**Figure S5. Evaluation of how approach and attachment to the anesthetized mother vary with task experience.**

**(a)** (left) Cumulative incidence curves showing the cumulative fraction of pups that approached the dam at each latency in each of the 8 ordered trials for non-ablated pups in **Fig 5b,c**. Each colored trace represents the cumulative incidence curve for one trial. (right) Change in hazard ratio per trial (tick) and 95% confidence interval (line) for the propensity to approach the dam (see Methods). **(b)** cumulative incidence curves showing the cumulative fraction of pups that approached the dam at each latency in each of the 8 ordered trials for DTA-ablated pups in **Fig 5b,c**. Conventions are as in **panel a.** **(c)** Cumulative incidence curves showing the cumulative fraction of pups that attached to the dam following a successful approach at each latency in each of the 8 ordered trials for non-ablated pups in **Fig 5b,c**. Conventions are as in **panel a.** **(d)** Cumulative incidence curves showing the cumulative fraction of pups that attached to the dam following a successful approach at each latency in each of the 8 ordered trials for DTA-ablated pups in **Fig 5b,c**. Conventions are as in **panel a.** (N.S. - not significant at $p=0.05$, **** - Significant at $p<0.0001$; multilevel mixed-effects Cox regression accounting for individual and litter effects N=23 ablated pups and 22 non-ablated pups in 6 litters; see Methods).

### SUPPLEMENTAL FIGURES

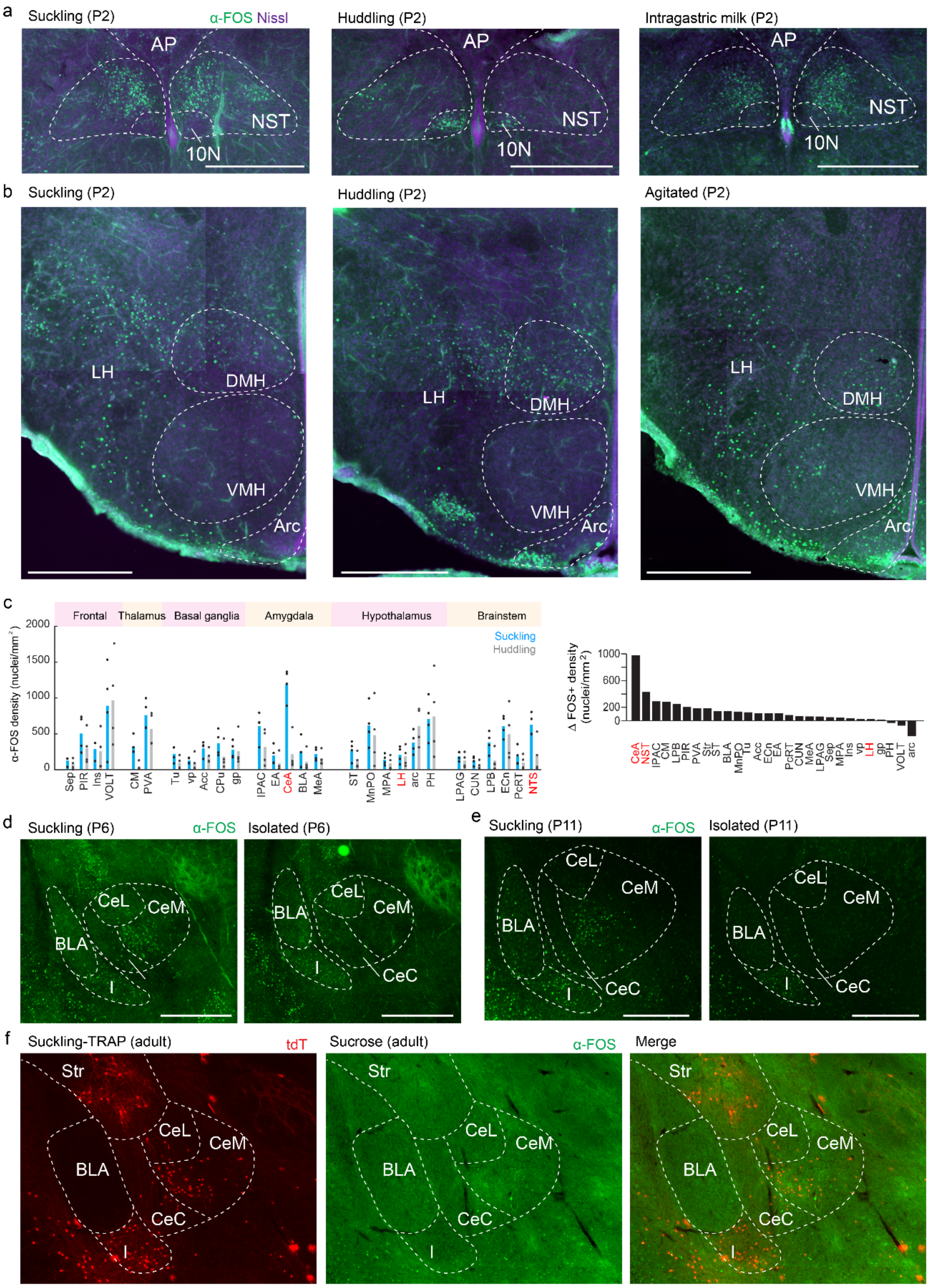


Figure S1

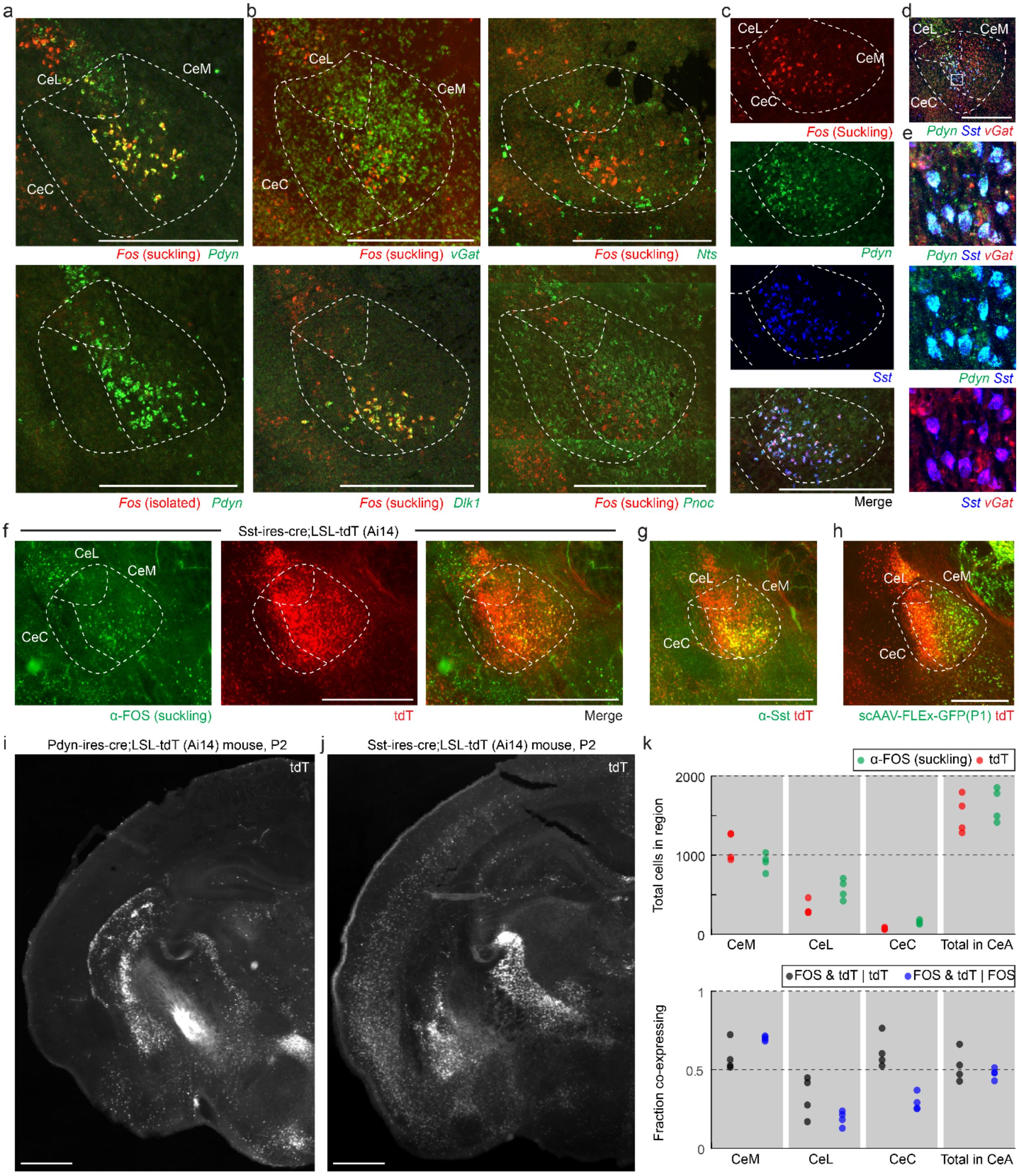
a
b
c
d
CeL
CeM
CeC
Fos (suckling) Pdyn
Fos (suckling) vGat
Fos (suckling) Nts
Fos (Suckling)
Pdyn Sst vGat
e
Pdyn
Pdyn Sst vGat
Fos (isolated) Pdyn
Fos (suckling) Dlk1
Fos (suckling) Pnoc
Sst
Pdyn Sst
Merge
Sst vGat
f
Sst-ires-cre;LSL-tdT (Ai14)
g
h
α-FOS (suckling)
tdT
Merge
α-Sst tdT
scAAV-FLEx-GFP(P1) tdT
i
Pdyn-ires-cre;LSL-tdT (Ai14) mouse, P2
j
Sst-ires-cre;LSL-tdT (Ai14) mouse, P2
k
α-FOS (suckling) tdT
2000
1000
0
Total cells in region
CeM
CeL
CeC
Total in CeA
FOS & tdT | tdT
FOS & tdT | FOS
1
0.5
0
Fraction co-expressing

Figure S2

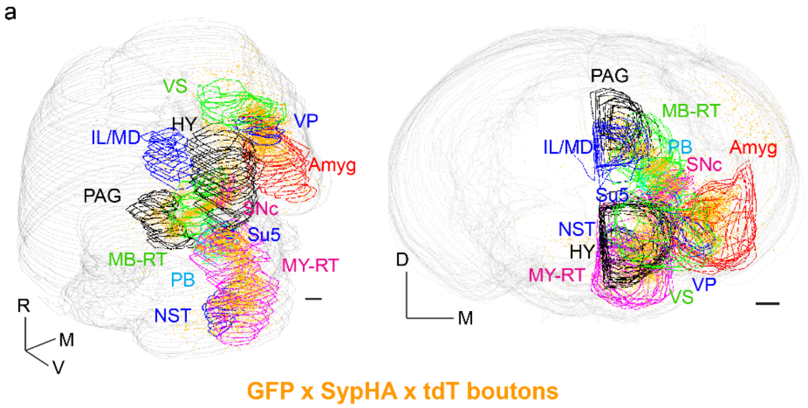
a
VS
VP
IL/MD
HY
Amyg
PAG
SNc
Su5
MB-RT
MY-RT
PB
NST
R
M
V
PAG
MB-RT
IL/MD
PB
Amyg
SNc
Su5
NST
HY
MY-RT
VP
VS
D
M
GFP x SypHA x tdT boutons

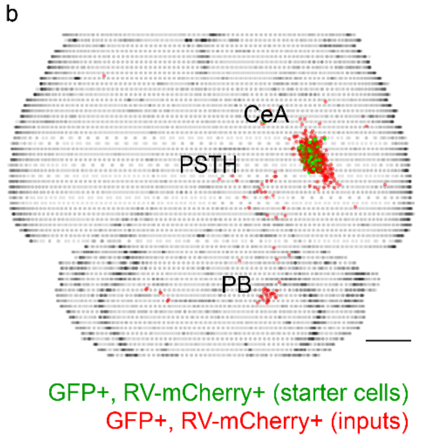
b
CeA
PSTH
PB
GFP+, RV-mCherry+ (starter cells)
GFP+, RV-mCherry+ (inputs)

Figure S3

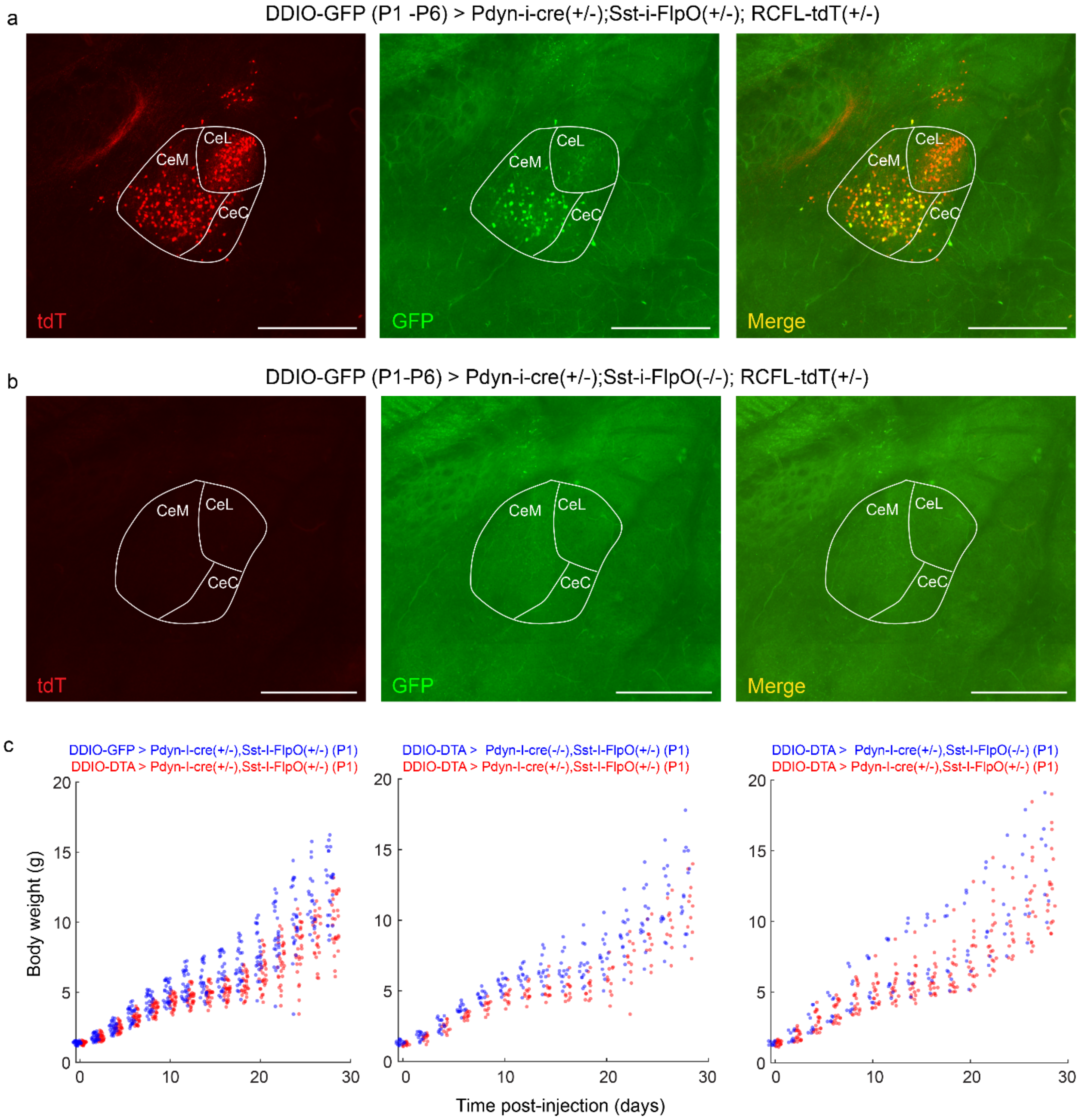
a
DDIO-GFP (P1 -P6) > Pdyn-i-cre(+/-);Sst-i-FlpO(+/-); RCFL-tdT(+/-)
CeL
CeM
CeC
tdT
GFP
Merge
b
DDIO-GFP (P1-P6) > Pdyn-i-cre(+/-);Sst-i-FlpO(-/-); RCFL-tdT(+/-)
c
DDIO-GFP > Pdyn-I-cre(+/-),Sst-I-FlpO(+/-) (P1)
DDIO-DTA > Pdyn-I-cre(+/-),Sst-I-FlpO(+/-) (P1)
DDIO-DTA > Pdyn-I-cre(-/-),Sst-I-FlpO(+/-) (P1)
DDIO-DTA > Pdyn-I-cre(+/-),Sst-I-FlpO(+/-) (P1)
DDIO-DTA > Pdyn-I-cre(+/-),Sst-I-FlpO(-/-) (P1)
DDIO-DTA > Pdyn-I-cre(+/-),Sst-I-FlpO(+/-) (P1)
Body weight (g)
Time post-injection (days)
0
5
10
15
20
30

Figure S4

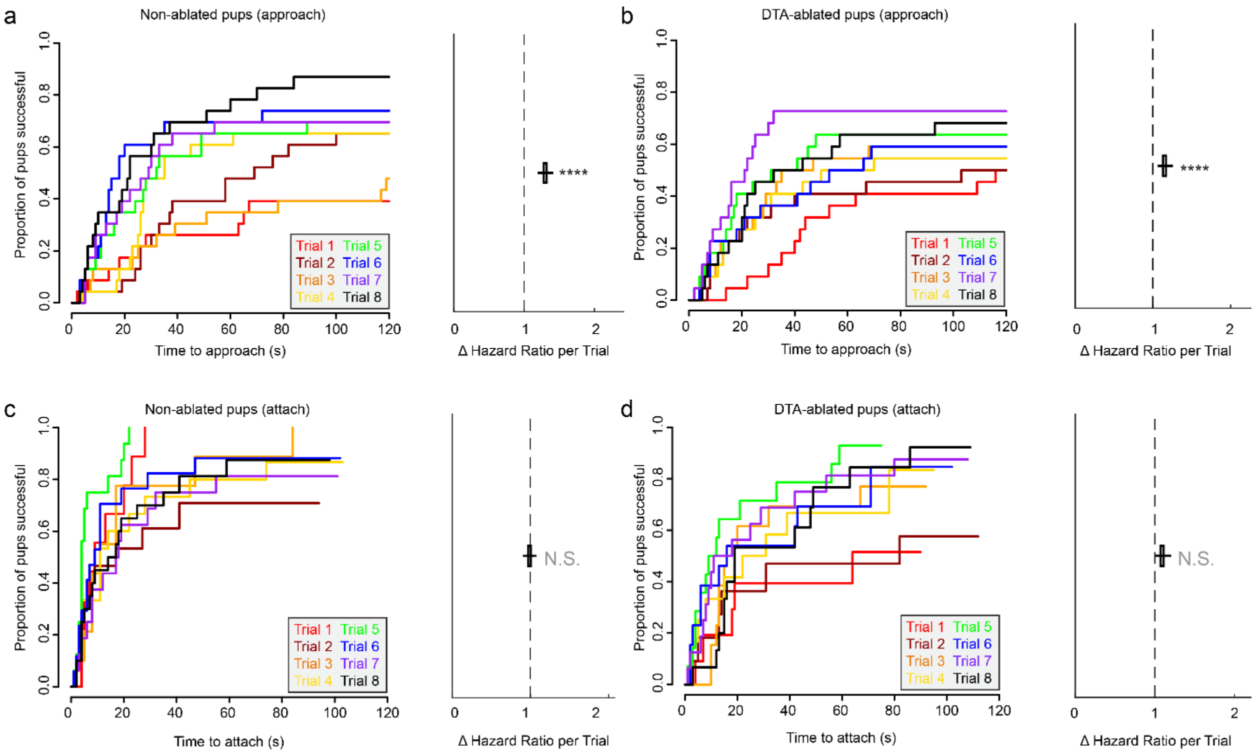
a
Non-ablated pups (approach)
Proportion of pups successful
Time to approach (s)
Δ Hazard Ratio per Trial
****
Trial 1 Trial 5
Trial 2 Trial 6
Trial 3 Trial 7
Trial 4 Trial 8
b
DTA-ablated pups (approach)
Proportion of pups successful
Time to approach (s)
Δ Hazard Ratio per Trial
****
c
Non-ablated pups (attach)
Proportion of pups successful
Time to attach (s)
Δ Hazard Ratio per Trial
N.S.
d
DTA-ablated pups (attach)
Proportion of pups successful
Time to attach (s)
Δ Hazard Ratio per Trial
N.S.

Figure S5
